# Homoploid hybrid speciation between cryptic species of a hidden snout weevil

**DOI:** 10.1101/2025.11.12.688067

**Authors:** Víctor Noguerales, Marius H. Eisele, Sophia L. Gray, Sean Stankowski, Brent C. Emerson

## Abstract

Hybridisation may generate a variety of evolutionary outcomes, including novel hybrid species without a change in ploidy. Within this process, referred to as homoploid hybrid speciation (HHS), selection on recombinant traits is often invoked as the driver of reproductive isolation (RI) between hybrid and parental species. However, our understanding of the temporal, geographic and phenotypic dimensions within which hybrid populations evolve RI against parental species is still limited. Here, we describe HHS within a beetle species complex that presents no obvious phenotypical or ecological differences among hybrid and parental species, and where all occur sympatrically. Demographic modelling supports hybrid speciation, where the hybrid lineage originated from historical admixture between parental lineages. Genome-wide estimates of heterozygosity and individual-based simulations of hybrid speciation provide further support for an ancient, rather than recent origin of the hybrid species. We leverage geographic patterns of genomic relatedness between hybrid and parental individuals to identify the likely origin within a hybrid swarm that was geographically isolated from both parental species. We argue that hybridisation is not just a driver of speciation, but may also act as a facilitator by enhancing the pathway to RI above that expected within a strictly allopatric model.

## Introduction

Homoploid hybrid speciation (HHS) appears to be relatively rare in nature, but evidence for it is increasingly being detected through analyses of genome-wide nuclear genetic variation that reveal mosaic genomes that are assigned to more than one parental species (e.g., Elgvin et al., 2017; Wang et al., 2022; Lopes et al., 2023). HHS, in its most basic definition, is the formation of new species that are reproductively isolated from the parental species from which they have been derived through hybridisation, and where ploidy is the same as that of the parental species (Eroukhmanoff et al., 2013; Nieto Feliner et al., 2017; Runemark et al., 2019). Homoploid hybrid lineages are increasingly being detected with genome-wide genetic markers, but direct assessment of reproductive isolation (RI) from both parental species is often limited by allopatric distributions. However, notable examples include the Italian sparrow (Passer italiae; Runemark et al., 2018) and cryptic species within the torpedo scad (Megalapsis cordyla; Muto et al., 2025).

Beyond the basic definition, there are divergent opinions about precisely what defines a hybrid species. Perhaps the most stringent definition is that of Schumer et al. (2014), who suggest that hybrid species should meet three criteria. The first is that there should be strong RI between parental and hybrid species. The second is that hybridisation should be evidenced by genetic data. The third is that RI should be directly derived from hybridisation itself. This definition seeks to identify hybridisation-derived isolating barriers, such as assortative mating, and ecological or genetic incompatibilities, as opposed to those that arise after hybridisation due to sorting of ancestral variants or new mutation (Schumer et al., 2018). In contrast, Nieto Feliner et al. (2017) have argued that by being overly restrictive, the definition of Schumer et al. (2014) may underestimate the importance of hybridisation in generating new species, because few species will satisfy all three criteria. This is particularly true for the third criterion, that RI should be derived from hybridisation itself, as it is difficult to test within hybrid lineages of older origin (e.g., Noguerales & Ortego, 2022; Zou et al., 2022), although in some cases this can be addressed with whole genome data when candidate genes associated with RI can be identified (Long & Rieseberg, 2024).

Because strong RI is typically polygenic, Long and Rieseberg (2024) argue that HHS may be more common and protracted than has been suggested by conceptual arguments and theoretical studies. While the criteria of Schumer et al. (2014) imply hybridisation itself to be important for the establishment of initial reproductive barriers against parental species, Long and Rieseberg (2024) propose an extended model. They point out that many models of hybrid speciation involve the sorting of parental species barriers, such that hybrid lineages will initially be less strongly isolated from the parental species than the parental species are to each other. Thus, as admixed populations form and diverge, even in the face of initially maladaptive or neutral admixture, they should at some point lead to HHS. This polygenic model is still consistent with the model of Schumer et al. (2014, 2015), but with protracted rather than instantaneous (or close to instantaneous) RI (Long & Rieseberg, 2024).

In this vein, Long and Rieseberg (2024) highlight the temporal dimensionality within which HHS might be expected to occur. Another important axis of variation that Long and Rieseberg (2024) describe is the extent of divergence between parental lineages at the time of a hybridisation event leading to hybrid origin, which by definition occurs within the so-called “grey zone of speciation” (i.e., where 0 < RI < 1; Roux et al., 2016). Hybrid speciation is necessarily set within a history of speciation with gene flow, with empirical support for a Goldilocks zone of divergence (Blanckaert & Bank, 2018; Corneault & Matute, 2018) that may be optimal for hybrid speciation to occur. The Goldilocks zone was originally described in the context of specific linkage architectures and intermediate recombination rates that might favour hybrid speciation (Blanckaert & Bank, 2018; see also Blanckaert et al., 2023), but has been also applied by Corneault and Matute (2018) to demonstrate that assortative mating is more pronounced within hybrid lineages at intermediate levels of divergence between hybridising parental species.

An area of interest toward a broader understanding of HHS is the relative roles of prezygotic, extrinsic postzygotic and intrinsic postzygotic isolation as facilitators of hybrid speciation. Given that hybrids are considered to likely be ecologically intermediate (Coyne & Orr, 2004), and that overlapping mating signals and preferences of hybrid populations with parental species are also likely to limit hybrid species formation, HHS has been considered to be infrequent (Schumer et al., 2014). However, in contrast, the emergence of transgressive phenotypes may favour the establishment of reproductive barriers between hybrid populations and their parental species (Dittirich-Reed & Fitzpatrick, 2013; Kagawa et al., 2023). With few exceptions, most examples of HHS involve parental species characterised by obvious ecologically or behaviourally relevant trait divergence (e.g., Schwarzbach et al., 2001; Barrera-Guzmán et al., 2018; Gompert et al., 2006; Lamichhaney et al., 2018; Schwarz et al., 2005), within which hybridisation may give rise to transgressive phenotypes.

Here we present a case of hybrid speciation in the absence of obvious ecological divergence between parental species, wherein the hybrid species and both parental species all occur sympatrically within a geographically limited area. *Silvacalles instabilis* (Wollaston, 1864) is a species of hidden snout weevil (Coleoptera: Curculionidae), endemic to the island of Tenerife on the Canary Islands (Figure 1), and restricted to the range-limited cloud forest formations within the island. Structured sampling has found *S. instabilis* to be limited to above-ground vegetation (Salces-Castellano et al., 2020) and associated with a variety of tree species (e.g., Laurus novocanariensis, Picconia excelsa, Persea indica, Myrica faya, Ilex platyphylla; Stüben et al., 2010), where it likely feeds on dead wood, as is characteristic of many species within the subfamily Cryptorhynchinae. Silvacalles instablis is one of 16 species of Coleoptera that have previously been characterised with genome-wide nuclear data across the cloud forest of the Anaga peninsula of Tenerife (Salces-Castellano et al., 2020). In the case of *S. instabilis*, in addition to the genome-wide nuclear data used in Salces-Castellano et al. (2020), several divergent nuclear genotypes, indicative of a probable second species, were originally excluded from analysis, and assumed to belong to the sister species *S. nubilosus* (Wollaston, 1864). However, a posterior revision of this material using a reference library of mitochondrial barcodes for the weevils of the Macaronesian region (Stüben, 2022) unequivocally assigns the divergent genotypes to *S. instabilis*. Here, we first clarify species boundaries and taxonomic assignment with a broader sampling of *S. instabilis*, together with individuals of *S. nubilosus*. We then assess population structuring of nuclear genomic variation within *S. instabilis*. We finally leverage the genome-wide nuclear data for a demographic modelling approach to understand the dynamics of gene flow and admixture within the speciation history of the *S. instabilis* complex. Our results provide unequivocal support for an initial cladogenetic event that underpins the origin of two species, followed by admixture between these two parental lineages giving rise to a third hybrid species. We discuss the well-documented spatio-temporal dynamics of species ranges within the focal habitat, and the potential for these to have catalysed hybrid speciation. We then discuss the significance of our results in the context of HHS theory.

**Figure 1.**
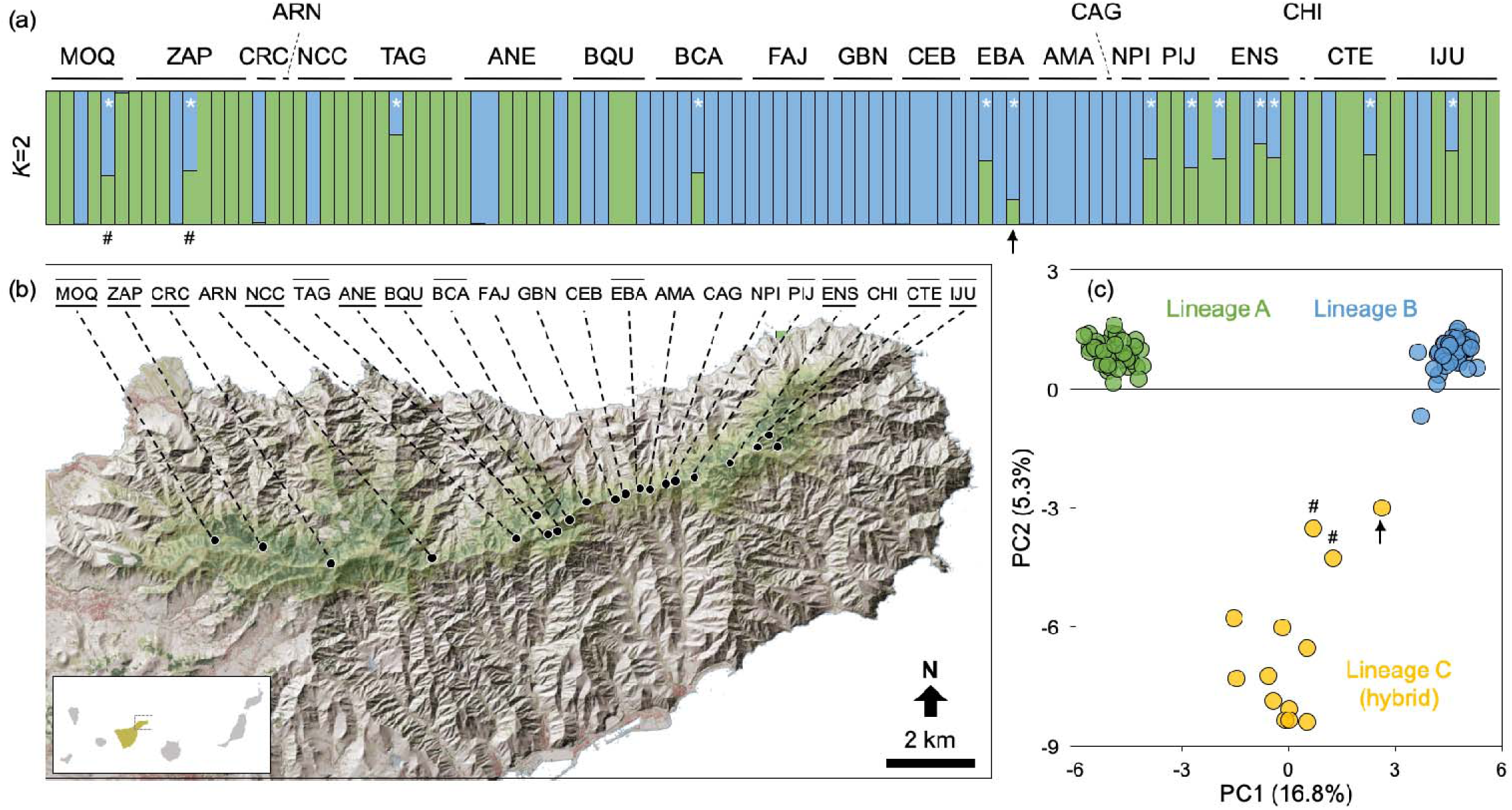
Panel (a) depicts the ancestry coefficients per individual within the *Silvacalles instabilis* species complex, as inferred in structure assuming two ancestral populations (*K*=2). Panel (b) shows the geographic location of each sampling site along the Anaga peninsula. Panel (c) shows a principal component analysis (PCA) of genetic variation for the sample individuals. In the upper plo, vertical bars with a white asterisk represent individuals of admixed co-ancestry, whose position within the PCA is represented by orange circles. Black arrows in panels (a) and (c) are used to highlight the *F*_1_ between lineages B and C, while hash symbols identify the two westernmost hybrid individuals. In panel (b), underlined sampling site codes represent those where single-ancestry individuals from lineages A and B co-occur. Site codes with a line above indicate where hybrid individuals co-occur with at least one parental lineage. Population codes as in Table S1.

## Materials and methods

### Sample collection

Previous sampling of *Silvacalles instabilis* by Salces-Castellano et al. (2020) was enhanced with 11 additional sites along the dorsal ridge of the Anaga peninsula (Noguerales & Emerson, 2025), yielding a total of 106 specimens from 21 sampling sites (Figure 1; Table S1). Four additional individuals were also collected from the sister species *S. nubilosus*, which were included as an outgroup in downstream genomic analyses when required. Supplementary Methods S1 provides further details on sampling.

### Mitochondrial and genome-wide nuclear data

Genomic DNA was extracted from each individual using the Biosprint DNA Blood Kit (Qiagen) on a Thermo KingFisher Flex automated extraction instrument. The barcode region of the mitochondrial DNA (mtDNA) cytochrome c oxidase subunit I (COI) was amplified for each individual using the primers Fol-degen-for and Fol-degen-rev (Yu et al., 2012), following conditions described in Noguerales and Emerson (2025). PCR products were purified with enzymes ExoI and rSAP (New England Biolabs, Ipswich, MA, USA), and then Sanger sequenced (Macrogen, Madrid, Spain), edited with geneious prime version 2021.1.1 and aligned using mafft (FFT-NS-i method; Katoh & Standley, 2013).

To generate genome-wide nuclear data, individual genomic DNA was processed using the double-digestion restriction-site associated DNA sequencing protocol (ddRADseq, Peterson et al., 2012), as described in Salces-Castellano et al. (2020), and Noguerales and Emerson (2025).

In brief, individual DNA extracts were digested with the restriction enzymes MseI and EcoRI (New England Biolabs). Resulting genomic libraries were then pooled at equimolar ratios, followed by size selection for fragments between 200 and 300 base pairs (bp), and paired-end sequenced (150 bp) on an Illumina NovaSeq6000 (Novogene, Cambridge, UK).

### Genetic relatedness using mtDNA data

Phylogenetic relationships among all individual mtDNA sequences were assessed with a maximum-likelihood approach as implemented in iq-tree version 1.6.12 (Trifinopoulos et al., 2016). A total of 1000 ultrafast bootstrapping replicates were conducted for node support calculation (Minh et al., 2013), implementing a HKY+I model of nucleotide evolution according to the Bayesian information Criterion (BIC), as calculated by modelfinder within iq-tree (Kalyaanamoorthy et al., 2017). Four individuals belonging to the sister species *Silvacalles nubilosus* were used as an outgroup.

#### De novo assembly of ddRADseq data

Raw sequences were demultiplexed, quality-filtered and de novo assembled using ipyrad version 0.9.96 (Eaton & Overcast, 2020). Briefly, quality-filtered reads were clustered considering a threshold of sequence similarity of 85% (clust_threshold), and clusters with a minimum coverage depth <5 (mindepth_majrule) and a maximum coverage depth >10,000 (maxdepth) were discarded. Statistical base calling was then performed at a minimum depth of 6 (mindepth_statistical). Finally, loci that were not present in at least 80% of individuals (min_saples_locus) were discarded. Unless otherwise indicated (see phylogenetic analyses in raxml), all downstream analyses were performed using datasets of unlinked SNPs (i.e., a single SNP per RAD locus). Supplementary Methods S2 provides further information on sequence assembly and data filtering in ipyrad.

### Genomic clustering analyses

Nuclear relatedness among individuals and potential signatures of admixture were analysed using the Bayesian clustering method implemented in structure version 2.3.3 (Pritchard et al., 2000). A hierarchical approach was used to identify the underlying genomic variation at different evolutionary scales (Noguerales et al., 2016; Ortego et al., 2021). Accordingly, (i) a global analysis was first conducted to evaluate overall population genetic structure across all individuals, and then (ii) analyses were repeated within inferred genetic groups to test for more fine-scale genetic structure within each. For all levels of analysis, two statistics were used to interpret the range of ancestral populations (K) that best describes the data: log probabilities of data (Pr(X|*K*); Pritchard et al., 2000) and Δ*K* (Evanno et al., 2005), both calculated in structure harvester version 0.7 (Earl & vonHoldt, 2012).

In addition to the Bayesian clustering analyses, major axes of genomic variation were visualised with a Principal Component Analysis (PCA) using the ‘gl.pcoa’ function as implemented in the package dartR version 2.9.7 (Mijangos et al., 2022) and r version 4.2.2 (R Core Team, 2022). As for structure, a hierarchical approach was followed, thus PCAs were conducted at different evolutionary scales, both across all individuals and within each of the resulting genetic groups. Following Ortego and Noguerales (2025), both structure and PCA analyses were performed with datasets only including SNPs with a minor allele frequency (MAF) >3%, using the ‘gl.filter.maf’ function as implemented in the r package dartR. Further details on genomic clustering analyses are provided in Supplementary Methods S3.

### Analysis of nuclear genomic diversity and admixture

Elevated heterozygosity is expected in hybrid individuals as a result of the admixture of parental forms with divergent genetic backgrounds and thus carrying different alleles for a given nucleotide site (Boca et al., 2020). Elevated heterozygosity will be highest in first-generation hybrid (*F*_1_) individuals and will diminish over time as a function of crossing dynamics and genetic drift (Fitzpatrick, 2012). Thus, the detection of elevated heterozygosity provides some indication of the recency of admixture. To assess heterozygosity, genome-wide nuclear data were used to estimate the observed heterozygosity (*H*_O_) per individual as implemented with the ‘gl.report.heterozygosity’ function of the r package dartR. This analysis was repeated across all individuals together and independently within the main lineages detected with structure and PCA.

Additionally, the presence of different hybrid classes (i.e., *F*_1_, F_2_, backcrosses and later-generation hybrids) was evaluated through estimating hybrid index (*Hb*_i_) and interclass heterozygosity (_i_*He*) per individual, as calculated with the r package triangulaR version 0.0.1 (Wiens et al., 2025). Individuals were assigned to putative parental and hybrid populations according to the main genetic groups inferred with clustering analyses (structure and PCA). For all analyses, a range of allele frequency thresholds was used (δ = [0.7, 0.8, 0.9]) to identify ancestry-informative markers (AIMs) across parental demes for the estimation of both *Hb*_i_ and _i_*He* values per individual.

### Phylogenomic inference

Evolutionary relationships at the levels of individuals, and the higher-order genetic groups detected with structure and PCA, were assessed through two complementary approaches using genome-wide nuclear information. First, phylogenetic relationships were reconstructed among all individuals using a maximum-likelihood approach as implemented in raxml version 8.2.12 (Stamatakis, 2014). Analyses were performed on a matrix of concatenated SNPs (i.e., including all SNPs per RAD locus), with ascertainment bias correction based on the conditional likelihood method (Lewis, 2001), using the GTR-GAMMA model of nucleotide evolution. The best-scoring maximum likelihood tree and nodal support were estimated through 100 rapid bootstrap replicates. Four individuals belonging to the sister species *Silvacalles nubilosus* were used as an outgroup (Table S1). Second, phylogenetic relationships among higher-order genetic groups were reconstructed via the Bayesian multispecies coalescent method implemented in snapp version 1.5.2 (Bryant et al., 2012), using two individual grouping strategies: (i) individuals were assigned to the main genetic groups identified within the global analyses of both structure and PCA, and; (ii) to take into account geographical structuring of genetic variation within the higher-order genetic groups, individuals were assigned into geographic populations of single co-ancestry, based on results of within-lineage analyses by both structure and PCA. Extended information on phylogenomic inference is provided in Supplementary Methods S4.

### Testing for introgression

Four-taxon ABBA-BABA tests based on D-statistics (Durand et al., 2011) were used to determine the role of gene flow between non-sister taxa in explaining genomic admixture patterns. This method enables the evaluation of the extent to which an excess of shared alleles in a given lineage is the result of either historical gene flow from a non-sister lineage, or retention of ancestral genetic variation (i.e., incomplete lineage sorting, ILS). Using the topology retrieved in raxml and snapp, where the sister lineages B and C have diverged from lineage A (Figure S1-S2), significantly positive or negative D-statistics would support the existence of gene flow between the non-sister lineages A and C or between A and B, respectively. The two grouping strategies used for the phylogenomic analyses in snapp used were also employed for ABBA-BABA tests. First, all individuals were assigned to lineages described within the global analyses of both structure and PCA. Second, structuring of genetic variation within the higher-order genetic groups (structure and PCA) was taken into account. Accordingly, individuals were grouped into geographic populations of single co-ancestry on the basis of the clustering schemes observed in the within-lineage analyses (structure and PCA) and then D-statistics were recalculated. For all analyses, the four individuals from *Silvacalles nubilosus* were used as an outgroup. ABBA-BABA tests were conducted in ipyrad using 1000 bootstrap replicates to obtain the standard deviation of the D-statistics. Extended information on introgression tests is provided in Supplementary Methods S5.

### Testing alternative demographic models

A coalescent simulation-based approach was applied to statistically evaluate the fit of the observed data to alternative scenarios of divergence and hybridisation (Figure S3). To this end, the composite likelihood of the observed data was estimated under a specified demographic model using the site frequency spectrum (SFS) and the simulation-based approach implemented in fastsimcoal version 2.5.2.21 (Excoffier et al., 2021). Six competing demographic scenarios were evaluated assuming: (i) strict isolation with hierarchical divergence among lineages (hereafter, ‘strict bifurcation model’); (ii) directional introgression among non-sister lineages on a strict bifurcation tree (‘introgression model’; Noguerales & Ortego, 2022); (iii) strict isolation after a single ancestral origin of all lineages (‘polytomy model’, Zhao et al., 2023); (iv) directional introgression from one of the two parental lineages to the third putative hybrid lineage on a polytomous tree (‘polytomy followed by introgression model’); (v) the same as (iv), but for the other parental lineage, and finally; (vi) fusion of the two parental lineages giving rise to a third putative hybrid lineage (‘hybrid speciation model’, Barrera-Guzmán et al., 2018; Noguerales & Ortego, 2022; Emerson et al., 2025). For demographic models i and ii, the topology inferred from phylogenomic analyses in raxml and snapp was used. Hybridisation events in models ii, iv, v and vi were parameterised using the γ parameter, which represents the expected proportion of migrants moving from a given source deme to a sink deme (Excoffier et al., 2013). These six models were evaluated either assuming contemporary symmetrical migration among all demes, or the absence of migration, thus yielding a total of twelve competing demographic scenarios (Figure S3). Details on composite likelihood estimation, model selection approach and calculation of confidence intervals for parameter estimates under the most-supported model are described in Supplementary Methods S6.

### Individual-based simulations of hybrid speciation

Given that the hybrid speciation model was best supported by demographic inference (see Results section), a custom simulation framework (S. L. Gray, https://github.com/sophiagray2001/hybridsimulator.git) was used to understand how rates of drift and migration might have shaped the genomic variation of the hybrid population, in terms of the ancestry (hybrid index, *Hb*_i_) and genetic diversity (interclass heterozygosity, _i_*He*) across individuals (see Supplementary Methods S7). The basic model consisted of two ancestral populations, each consisting of 100 individuals, that carry 58 semi-diagnostic loci with allele frequencies and rates of missing data that match the empirical dataset of AIMs for triangulaR analysis (i.e., loci with δ ≥0.8). These populations mate to form a population of 200 *F*_1_ hybrids. The hybrid population is then allowed to evolve for 3000 generations (equivalent to 15*N*_e_ generations), with each mating pair leaving 2 offspring.

A total of 13 variations on this basic model (50 independent replicates each) were explored. First, because the genomic distribution of loci is currently unknown, how the degree of linkage affects the trajectory of the hybrid populations was explored through simulating the joint distribution of *Hb*_i_ and _i_*He* under five alternative scenarios. These scenarios assumed: (i) free recombination among loci; (ii) the random assignment of loci to 20 linkage groups (the average chromosome number across species of Coleoptera) with rates of recombination between neighboring loci defined by their map distance (each replicate with a different randomly generated map); (iii) all markers assigned to a single chromosome; (iv) tight linkage where all loci were clustered within 0.5 centimorgans (cM) of one another, and; (v) clustering with no recombination. Following this, eight additional scenarios were built to explore the effect of incomplete RI. These scenarios allowed either gene flow from one parental population (scenarios vi-ix) or both parental populations (scenarios x-xiii) into the hybrid population, with each assuming four rates of migration (*m*=0.005, *m*=0.001, *m*=0.0005 and *m*=0.0001).

## Results

### Mitochondrial data and phylogenetic relatedness

A total of 105 mtDNA sequences were obtained, corresponding to 602 bp within the COI region. Maximum uncorrected genetic divergence between *Silvacalles instabilis* and *S. nubilosus* was 2.5%, with a maximum divergence of 1.8% within *S. instabilis*. Accordingly, iq-tree inferences revealed the existence of two well-supported monophyletic clades, corresponding to *S. instabilis* and *S. nubilosus* (Figure S4). Haplotypes within *S. instabilis* segregated into two major clades which did not reflect any clear phylogeographic structure or an association with the genetic groups inferred using genome-wide nuclear data (see further results). When mapped onto nuclear-based clustering inferences, these mtDNA clades were revealed to conform to polyphyletic groups of different nuclear ancestry (Figure S4).

### Genome-wide nuclear data

A total of 241.17 million raw sequences across the 110 individuals were retrieved: 106 individuals of *Silvacalles instabilis* and 4 specimens belonging to the sister species *S. nubilosus* (Figure S5). After the filtering steps in ipyrad, each individual retained an average of 2.19 million reads (SD=1.85 million, Figure S5a) and 40,671 clusters (SD=14,371; Figure S5b) of high-quality sequences, with a mean depth per locus of 33.01 (SD=27.08).

### Genomic clustering analyses

Global analysis across all individuals in structure identified the most likely number of ancestral populations to be *K*=2, under the Δ*K* criterion (Figure S6a). Individuals showing high single ancestry for one of the two ancestral populations were more commonly sampled in the western and eastern margins of the Anaga peninsula (hereafter ‘lineage A’), with individuals assigned to the other lineage being more commonly sampled more centrally within the peninsula (hereafter ‘lineage B’), between sampling sites BCA and NPI (Figure 1ab). However, single-ancestry individuals from both lineages were found sympatrically in 9 out of the 21 sampling sites. In addition, 13 individuals with strong signatures of genetic admixture (hereafter ‘lineage C’) were sampled at 9 sites, together with individuals of high single ancestry to either one or both ancestral populations (Figure 1ab). These admixed individuals showed mostly even co-ancestry coefficients (0.33 <q-value <0.67), with the exception of one individual from sampling site EBA, which exhibited higher ancestry from lineage B (q-value=0.81; Figure 1ac). In addition to the Δ*K* criterion, the log probability of data steadily increased until reaching an asymptote at *K*=3, indicating that genomic variation is hierarchically structured (Janes et al., 2017; Figure S6a). Accordingly, clustering with *K*=3 grouped admixed individuals into a third ancestral population of mostly single ancestry (Figure S1), in line with global PCA inferences (Figure 1c). The PCA across all individuals revealed that genomic variation is largely organised into three well-defined genetic groups (Figure 1c). Individuals showing an intermediate position between the two more divergent groups along the first component corresponded to those individuals showing an admixed genetic background (lineage C) for *K*=2 in structure (Figure 1a).

structure analyses restricted to either lineages A or lineage B revealed genomic variation to be structured into two geographically coherent ancestral populations within each, according to Δ*K* criterion (Figure S6bc). Within each lineage, individuals of single ancestry were sampled exclusively in either westernmost or easternmost sites. Additionally, within each lineage, admixed individuals were spatially intermediate between eastern and western populations (Figures S7-S8). Admixed individuals within lineage A were broadly distributed in central Anaga, between the sites of ARN and BQU, while lineage B exhibited a more abrupt change of co-ancestry, which coincided with the unique sampling site of FAJ. In agreement with structure inferences, lineage-specific PCAs described genomic variation within lineages A and B to be largely organised into west and east genetic groups within each (Figures S7-S8). Differences in allele frequencies within each lineage were well represented along the first component, with admixed individuals sampled in central Anaga having an intermediate position in their respective PCA, in concordance with the inferred geographic co-ancestry in structure (Figure S7-S8).

The log probability of the data for the within-lineage structure analyses increased beyond *K*=2 and stabilised at *K*=3 (Figure S6bc). When considering *K*=3 in lineage A, individuals of admixed ancestry from the central sites of ARN to BQU clustered into a third ancestral population of mostly single ancestry (Figure S7). Within lineage B, *K*=3 defined further geographic structuring within eastern Anaga, with individuals from the easternmost sampling sites of ENS, CHI, CTE and IJU assigning with high ancestry to a third ancestral population (Figure S8).

### Analysis of nuclear genomic diversity and admixture

Levels of observed heterozygosity (*H*_O_) across individuals ranged from 0.029 to 0.074 (Figure S1), with lineage C (admixed ancestry) presenting significantly lower *H*_O_ than that of lineages A and B) (ANOVA; F_2,103_=12.28; p-value <0.001; Figure S9). According to hybrid identification analyses with triangulaR (Figure 2), individuals within lineage C exhibited hybrid indices (*Hb*_i_) ranging from 0.38 to 0.70, with interclass heterozygosities (_i_*He*) as low as those observed within lineages A and B (Figure 2a). This pattern did not align with expectations for early-generation hybrids (Wiens et al., 2025), but rather with a much older origin of admixture. However, one individual assigned to lineage C presented a notably higher heterozygosity (_i_*He*=0.35) compared to parentals, or other admixed individuals, and a skewed co-ancestry (*Hb*_i_=0.76) toward lineage B which was in line with its structure-derived ancestry coefficient, suggesting it may represent a relatively recent hybrid between lineages B and C. Accordingly, when compared directly against putative parental lineages B and C, this individual exhibited an elevated heterozygosity (_i_*He*=0.61) and a co-ancestry value close to 0.5 (*Hb*_i_=0.41) (Figure 2b), consistent with expectations for an *F*_1_ hybrid. A comparison involving lineages A and C as putative parentals revealed this hybrid individual to be more closely related to lineage C (*Hb*_i_=0.71), and with relatively low heterozygosity (_i_*He*=0.30) (Figure 2c). Alternative allele frequency thresholds (δ=[0.7, 0.8, 0.9]) provided qualitatively similar results. Consequently, this individual was inferred to be a *F*_1_ hybrid between lineages B and C, and was discarded from further analyses.

**Figure 2.**
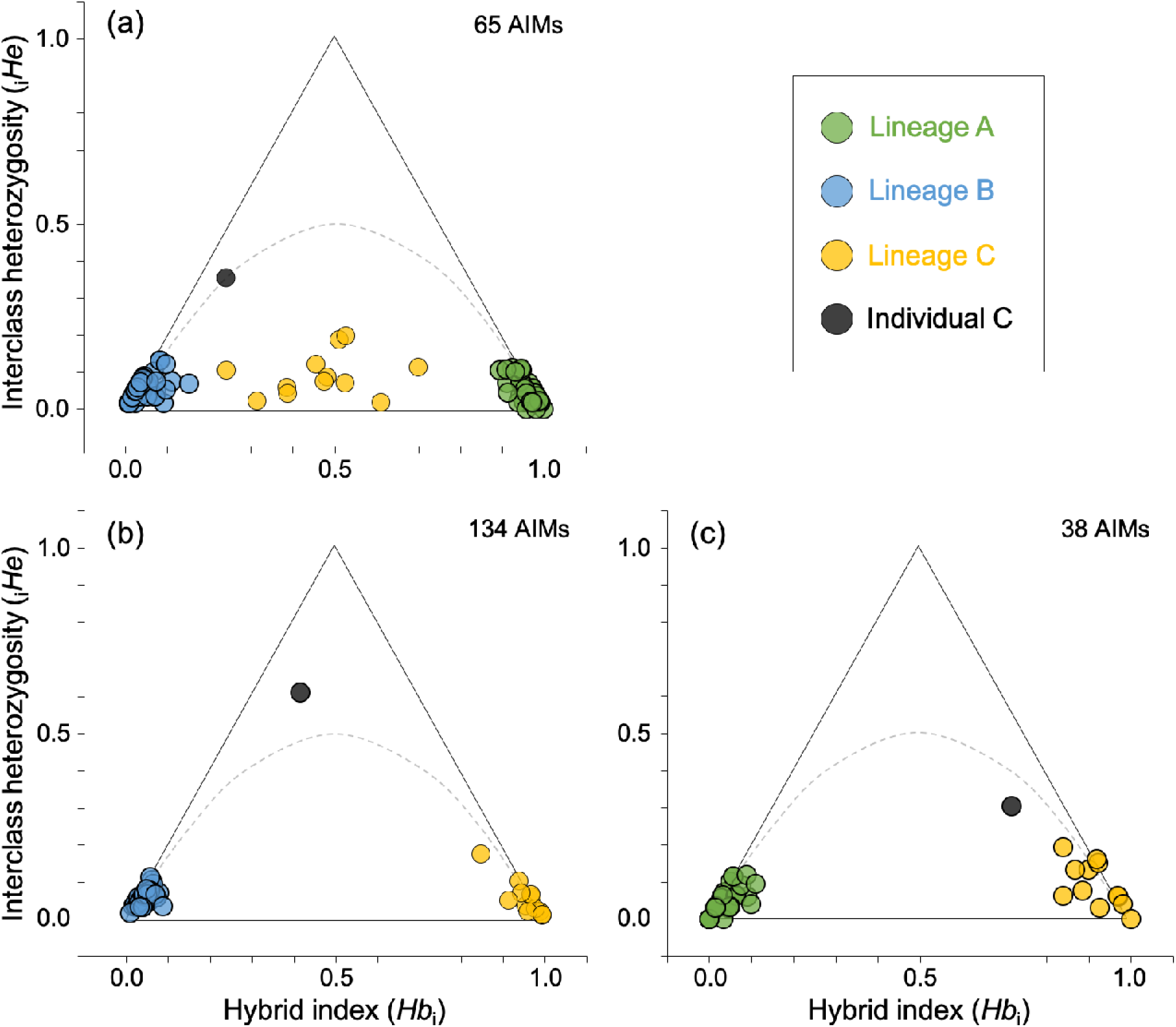
Triangular plots showing potential hybrid and putative parental lineages within the *Silvacalles instabilis* species complex, based on ancestry-informative markers (AIMs) identified using an allele frequency difference threshold of δ=0.8. Individuals from each lineage are represented by different colours. The black circle identifies an individual from lineage C that i identified as an *F*_1_ between lineages C and B.

### Phylogenomic inference

The phylogenetic tree reconstructed in raxml using all individuals revealed three well-supported monophyletic clades, consistent with the three lineages retrieved from the global structure and PCA analyses (Figure S1). The raxml topology supported a sister relationship between lineage B and lineage C (individuals of admixed co-ancestry), with earlier divergence giving rise to lineage A (Figure S1).

Coalescent reconstructions in snapp for the three lineages derived from previous genomic clustering (structure and PCA) provided similar topological inferences to the individual-based raxml tree. Accordingly, snapp supported an early split that gave rise to lineage A, followed by divergence between lineages B and C (Figure S2a). Similarly, snapp analyses accounting for geographic populations of single ancestry within lineages A and B revealed a similar topology (Figure S2b), supporting a sister relationship between lineages B and C. These analyses described a shallower divergence between the west and east groups within lineage B than that observed within their homologous geographic populations within lineage A (Figure S2b).

### Introgression and alternative demographic models

ABBA-BABA tests including all individuals from lineages A and B revealed highly significant introgression between the non-sister lineages A and C (Table S2). Significant introgression was also detected between either west or east lineage A and lineage C. This pattern remained significant regardless of whether the west or east groups from lineages A and B were considered within the topology, suggesting an introgression event predating the geographical structuring within lineage A and B (Table S2).

Coalescent demographic modelling in fastsimcoal identified the ‘hybrid speciation’ scenario (model vi) incorporating contemporary migration as the most-supported model (Table S3; Figure S3). Support for the next best model, the ‘introgression model’ (model ii) was substantially less with ΔAIC > 20. Among the remaining models, the scenario assuming a strict bifurcation divergence was the least supported (Table S3). The subset of models assuming no contemporary migration were much more poorly supported than their homologous scenarios considering contemporary migration (ΔAIC > 480; Table S3, Figure S3). Considering a 1-year generation time, parameter estimation in fastsimcoal within the most-supported model suggested parental lineages A and B diverged from a common ancestor approximately 0.83 Ma (T_DIV2_; 95% confidence interval [CI]: 0.65-0.97 Ma; Figure 3). The origin of lineage C as a result of hybridisation between parental lineages A and B was estimated to have taken place approximately 0.20 Ma (T_HYB_; CI: 0.16-0.24 Ma). Consistent with co-ancestry coefficients derived from structure, the γ parameter estimate was 0.60 (γ_HYB_; CI: 0.50-0.69), suggesting a relatively even contribution of parental lineages A and B to the gene pool of lineage C (Figure 1). Contemporary migration was estimated to be consistently very low across demes (Figure 3). However, the migration rate between the two parental lineages was estimated to be only slightly lower (m =7.90×10^-7^) than that between lineage C and either parental lineage A or B (m =1.05×10^-6^; m =1.13×10^-6^) (Figure 3).

**Figure 3.**
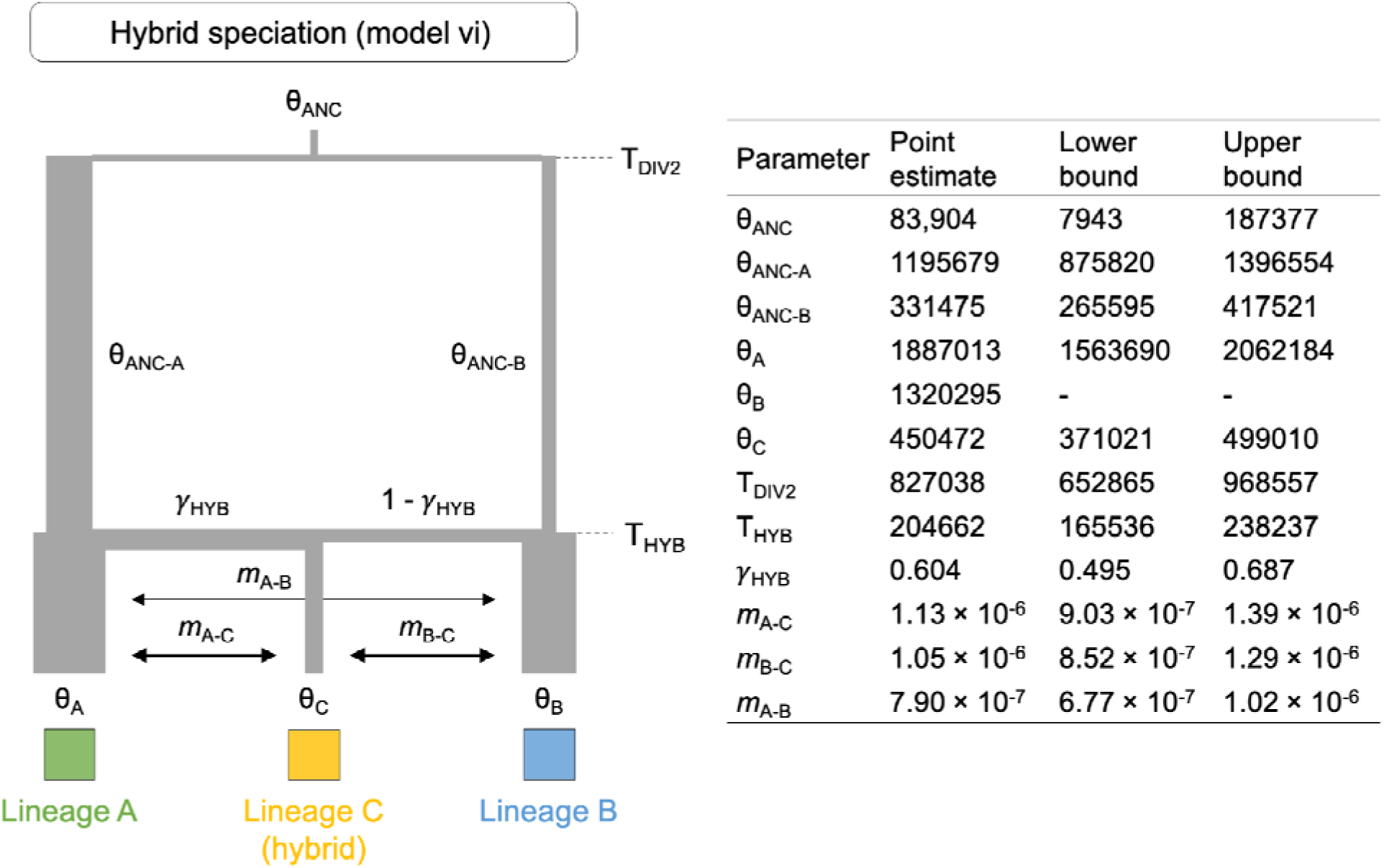
Parameters inferred from the coalescent simulations with fastsimcoal under the most-supported demographic model (Table S3). Point estimates and lower and upper 95% confidence intervals are shown for each parameter of the hybrid speciation scenario (model vi). Model parameters include ancestral (θ_ANC_, θ_ANC-A_, θ_ANC-B_) and contemporary (θ_A_, θ_B_, θ_C_) effective population sizes, timing of divergence (T_DIV2_) and hybridization (T_HYB_), contribution coefficient (γ_HYB_) and symmetrical migration rates per generation (*m*_I-J_). The parameters T_DIV2_ and T_HYB_ are expressed in years considering 1-year generation time. The relative magnitude of θ, T_DIV2_, T_HY_ and γ_HYB_ are represented by varying width and length of branches, while arrow thicknes indicates the relative magnitude of contemporary migration among demes (*m*_I-J_). Note that θ for lineage B (θ_B_) was previously calculated and fixed in fastsimcoal to enable the estimation of other parameters.

### Individual-based simulation of hybrid speciation

Individual-based simulations of hybrid speciation showed that linkage relationships of ancestry informative markers could influence both the rate of loss of heterozygosity (_i_*He*) due to drift, and the distribution of ancestry (*Hb*_i_) in hybrid populations. In simulations with free recombination (r=0.5) or loose linkage, similar rates of _i_*He* loss were observed, with the hybrid population (lineage C) reaching mean levels of heterozygosity of parental lineages A and B (mean _i_*He*_AB_) at around ∼7.35*N*_e_ generations (Figure 4ab; Figure S10). Although replicate simulations showed some variability in their ancestry scores, the distribution was tightly centred around 0.5 (Figure 4c). With tight linkage (all markers clustered within 0.5 cM), the rate of _i_*He* loss was far more variable, reflecting higher stochasticity due to the non-independence of linked sites (Figure S10). Similarly, the limited recombination of haplotypes led to a bimodal distribution of *Hb*_i_ scores, as the populations tended to fix one of the two ancestral haplotypes (Figure 4c; Figure S10).

**Figure 4.**
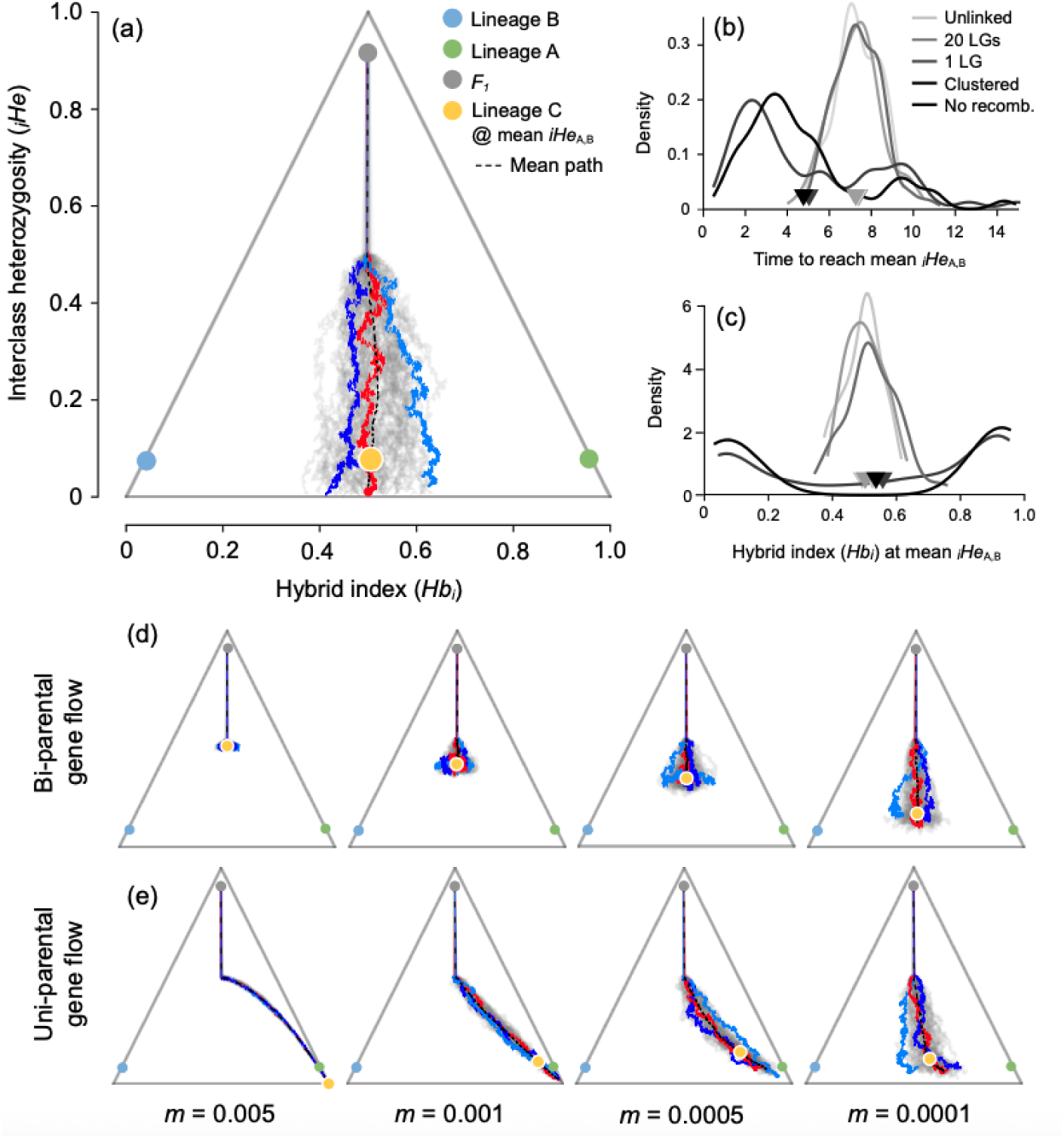
Individual-based simulations of hybrid speciation. (a) Triangle plot showing 50 replicate trajectories under a scenario where the 58 semi-diagnostic markers were arrayed on 20 linkage groups (LG). Each grey line represents the trajectory of one hybrid population ove time. The three highlighted trajectories in dark blue, light blue and red) represent a random path, an outlier, and the path closest to the overall mean, respectively. The black dashed line indicates the mean trajectory across all 50 replicates. The mean value of _i_*He* and *Hb*_i_ are shown for lineages A and B, and the *F*_1_ generation. The mean of lineage C at the time of reaching mean parental _i_*He* (mean _i_*He*_AB_) is indicated by the yellow circle. (b) Density plot showing the time (in *N*_e_ generations) to reach _i_*He*_AB_ across five scenarios with varying degrees of linkage. The x-axis shows time (*N*_e_ generations), and the y-axis shows the density of replicate simulations. (c) Density plot of ancestry proportions for each replicate at the time it reached mean _i_*He*_AB_ for the same linkage scenarios, with ancestry proportion on the x-axis and density on the y-axis. (d) Triangle plots of bi-parental migration (one migrant from each lineage A and B per event) under the free recombination scenario with migration rates of *m*=0.005, 0.001, 0.0005 and 0.0001, with colour and axis schemes identical to (a). (e) Triangle plots of uni-parental migration (one migrant from Lineage A) at the same four migration rates, with colour and axis schemes identical to (a).

Rates of migration from parental populations also had a strong effect on the trajectory of the hybrid populations. High migration from both parental populations (*m*=0.005) completely counteracted the loss of _i_*He* loss due to drift, such that the hybrid population remained highly heterozygous even after 15*N*_e_ generations (Figure 4d). With very low migration (*m*=0.0001), trajectories closely resembled complete RI, but with a slower rate of loss of _i_*He*. High and intermediate levels of migration from one parental produced asymmetrical trajectories, leading the hybrid population to become swamped by the influx of parental ancestry (Figure 4e). With lower migration, the bias in ancestry was far less pronounced, making the presence of weak gene flow harder to infer.

These simulations indicate that the AIMs identified in the *S. instabilis* complex are loosely linked, rather than being clustered in a genomic ‘island’ of differentiation, and that the hybrid population is strongly or fully isolated from both parental lineages. Here it is instructive to quantify the reduction in gene flow in terms of the strength of RI that would be needed to reduce mating to a given migration rate. For example, if we assume an equal probability of heterospecific (between lineage C and either lineage A or B) and homospecific encounters (within lineage C), then *m*=0.0001 is equivalent to an estimate of RI_4_=0.9998, according to Sobel and Chen (2014). These simulations were consistent with an old hybrid population, founded approximately ∼7*N*_e_ generations ago (95% CI: ∼5-9 *N*_e_ generations). However, this assumes that the loss of heterozygosity was due entirely due to drift. Any selection against heterozygotes would cause heterozygosity to be purged from the hybrid population at a much faster rate than by drift alone. Thus, the above value should be considered as an upper estimate.

## Discussion

Our results reveal that genetic variation for both the nuclear and mitochondrial genomes of *Silvacalles instabilis* is monophyletic with regard to *S. nubilosus*. This confirms the evolutionary independence of both taxonomic lineages, and is consistent with their classification into different species. However, within *S. instabilis*, genome-wide nuclear variation clearly segregates into three genetic groups, all of which are broadly sympatric across the cloud forest of the Anaga peninsula within the island of Tenerife. The maintenance of genotype differentiation in sympatry provides direct evidence that each of the three genetic groups is a biological species. Demographic modelling with genome-wide nuclear variation provides unequivocal support for an initial cladogenetic event that underpins the origin of two species, *S. instabilis* A and *S. instabilis* B. The subsequent evolution of the third species, *S. instabilis* C, was not a cladogenetic event. Rather, it is the product of admixture involving *S. instabilis* A and B.

### Hybrid speciation theory and *Silvacalles instabilis*

*Silvacalles instabilis* C is derived from hybridisation between *S. instabilis* A and B, supported by coalescent demographic simulations. Additionally, the sympatric distribution of the hybrid lineage with both parental species provides direct evidence for the RI of *S. instabilis* C from both parental species. *Silvacalles instabilis* C thus satisfies the first two of the three conditions for a hybrid species proposed by Schumer et al. (2014). What remains less clear is the extent to which RI may have been driven by hybridisation itself, the third and controversial criterion (e.g., Nieto Feliner et al., 2017) proposed by Schumer et al. (2014) for the definition of hybrid species.

Evidence for hybridisation as a driver of speciation can be found in hybridisation-derived phenotypes or incompatibilities that have acted as agents of isolation, which should not be confounded with those that are subsequent byproducts of admixture itself (Schumer et al., 2018). This criterion has focused attention on the evolution of RI in early-generation hybrids, with potential examples including Heliconius and Lycaeides butterflies (Gompert et al., 2006; Mavárez et al., 2006; Melo et al., 2009), Ragholetis fruitflies (Schwarz et al., 2005), manakin birds (Barrera-Guzmán et al., 2018), Darwin’s finches (Lamichhaney et al., 2018) and cichlid fishes (Olave et al., 2022), among others. However, Long and Rieseberg (2024) have highlighted the longer temporal dimension within which HHS might be expected to occur within a polygenic model. Incorporating the logic of Long and Rieseberg (2024) provides for a more holistic framework for characterising HHS, allowing for HHS to be a protracted process.

Recognised cases of HHS are typically associated with functional trait variation between parental species, where positive selection for intermediate or transgressive phenotypes simultaneously contributes to the RI of recombinant hybrid lineages (e.g., Nolte et al., 2005; Gompert et al., 2006; Meyer et al., 2006; Hermansen et al., 2011; Nice et al., 2013). Novel recombinant traits may also favour the colonisation of a novel habitat by an incipient hybrid species, thus facilitating establishment and early-stage persistence by allowing it to avoid introgression and competition with the parental species (Gompert et al., 2006). Consistent with this, and based on 28 studies of homoploid hybrid species of plants, Kadereit (2015) observed latitudinal/longitudinal and/or altitudinal ecogeographical displacement of hybrid lineages from parental lineages in the majority of cases. Within this framework of strong directional selection for intermediate or transgressive traits, evolution should act relatively quickly to drive speciation, analogous to a punctuated model of speciation, as has occurred with allopolyploidy diversification in plants (e.g., Linder & Rieseberg, 2004). In this context, *Silvacalles instabilis* presents an unusual case due to its extensive sympatry among species A, B and C, their shared ecology, and the absence of obvious phenotypical or ecological differences among the three species. Given that the existence of subtle variation among species cannot be excluded, the evolution of *S. instabilis* C through positive selection for intermediate or transgressive variants cannot be discounted. However, a plausible pathway to speciation in the absence of intermediate or transgressive phenotypes is the fixation of incompatibility loci from parental species (Schumer et al., 2015; Blanckaert & Bank, 2018).

### Selection against genetic incompatibilities and hybrid speciation

The conditions for hybrid speciation are enhanced when a hybrid population maintains a mixed combination of parental genetic incompatibilities that contribute to RI with both parental species. Theoretical work has demonstrated how reciprocal sorting of Bateson-Dobzhansky-Muller (BDM) genetic incompatibilities (Bateson, 1909; Dobzhansky, 1936; Muller, 1942) from parental species can result in hybrid speciation, even in cases where hybrids are selected against. Using simulations, Schumer et al. (2015) have shown that this is more likely for coevolving or adaptive incompatibility loci, where hybrid speciation can occur even in the face of ongoing gene flow from parental species and strong selection against hybrids. However, Blanckaert and Bank (2018), again using simulations, have demonstrated that hybrid speciation can also occur when parental BDM loci are evolving neutrally, and that speciation becomes highly probable under certain conditions. Within their neutral BDM model, Blanckaert and Bank (2018) highlight an apparent paradox. Barriers need to be weak enough to allow the formation of the hybrid population in the first place. However, at the same time barriers need to be strong enough for the hybrid population to be isolated from both parental populations. This paradox would be resolved within a dynamic where genomic reproductive isolating barriers between parental species are weak enough to allow a hybrid population to form, but with a subsequent period of geographic isolation of the hybrid population from parental species. Genomic signatures within *Silvacalles instabilis* are consistent with a geographically localised hybrid origin, and this has occurred within a topoclimatic arena within which isolation among populations is favoured during glacial periods (Salces-Castellano et al., 2020, 2021).

### Western origin and eastward range expansion of the hybrid species

A striking feature of genomic variation within *Silvacalles instabilis* C is the extent of genotypic differentiation among individuals, compared with the more limited differentiation seen within both parental species (Figure 1). Increased genotype diversity relative to parental populations is an expected outcome within early-generation hybrid crosses and backcrosses, as recombinant genotypes emerge from divergent parental allelic variation. However, this explanation can be discounted within *S. instabilis* C, as individuals are not early-generation hybrid crosses and backcrosses, as evidenced by the absence of elevated heterozygosity relative to the parental species (Figure 2). In contrast, a consideration of geographic patterns of individual relatedness both within and among species provides some explanation for the origin and significance of genotypic diversity within *S. instabilis* C.

Together with the *F*_1_individual between *Silvacalles instabilis* B and C, the two westernmost individuals of *S. instabilis* C (sampled from MOQ and ZAP, Figure 1b) present displacement in PCA space toward *S. instabilis* B (Figure 1c). The relatedness patterns for these two individuals may be explained by two competing scenarios. Within scenario one, subsequent to the origin of *S. instabilis* C, geographically localised introgression within the west of Anaga may have occurred from *S. instabilis* B to C. Within scenario two, the western extreme of Anaga may represent the geographic origin of *S. instabilis* C within Anaga, with subsequent range expansion to the east. Within this second scenario, higher sharing of ancestral variation would be expected between western populations of *S. instabilis* C and both parental species, with drift effects leading to increasing levels of differentiation of newly founded populations of *S. instabilis* C and both parental species, as range expansion of *S. instabilis* C occurs to the east. Both scenarios make testable predictions for geographic patterns of relatedness both within and among species.

Scenario one predicts higher relatedness of western individuals of *Silvacalles instabilis* C to *S. instabilis* B, but not to *S. instabilis* A. In contrast, scenario two predicts higher relatedness of western individuals of *S. instabilis* C to both parental species. Support for scenario two is provided by PCA analyses where only one of the parental species is included together with *S. instabilis* C (Figure 5). In both PCAs, both western individuals of *S. instabilis* C are displaced toward the parental species. In both cases, they are specifically displaced toward parental individuals from western sampling sites, consistent with an origin of admixture in the west. Further to this, in both PCAs *S. instabilis* C presents a geographic gradient of displacement from parental variation, with higher displacement associated with the more eastern sites of ENS, CTE and IJU. Results are thus consistent with a geographically localised western hybrid origin for *S. instabilis* C. Understanding how this western admixed population could have remained geographically isolated for some period from more eastern populations of the parental species requires consideration of how Quaternary climate oscillations have driven population isolation and secondary contact within dispersal-limited insect species of the Anaga cloud forest (Salces-Castellano et al., 2020, 2021).

**Figure 5.**
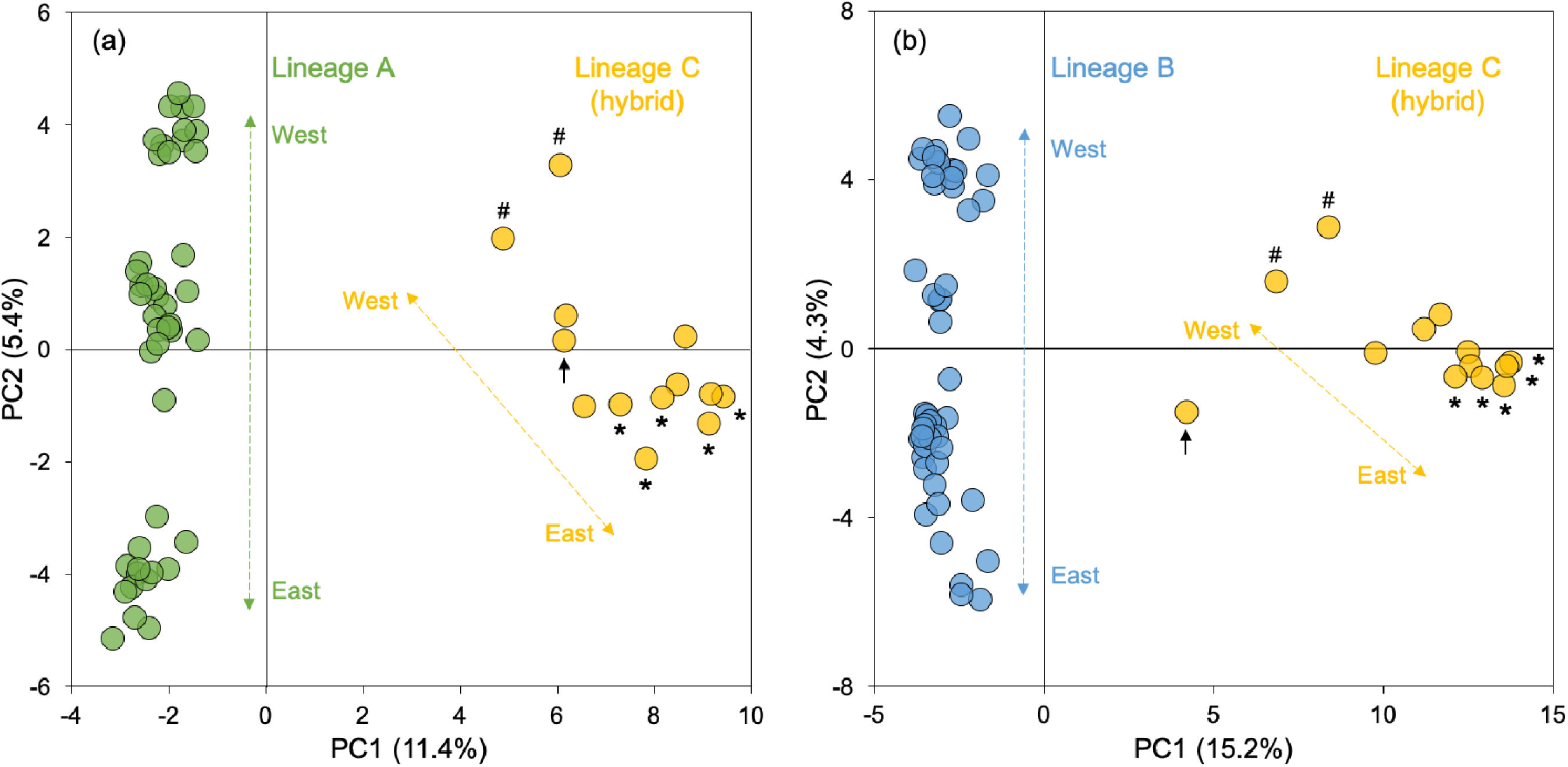
Structuring of genomic variation within hybrid lineage C with regard to one of the two parental lineages A and B, as inferred with a principal component analysis (PCA). Dashed and arrowed lines highlight west-east geographical structuring of genetic variation within lineages. For lineage C, individuals sampled in the westernmost sites (MOQ, ZAP) are highlighted with a hash symbol, with those sampled in the easternmost sites (ENS, CTE and IJU) being highlighted with an asterisk. As in Figure 1, the position of the *F*_1_ individual between lineages B and C (EBA site) is identified with a black arrow. Colours as in Figure 1.

### Isolation of hybrid population from parental species

Cloud forest connectivity and fragmentation within the Anaga peninsula has been moderated by climate oscillations throughout the Quaternary. Extensive evidence for this dynamic is provided by isolation and admixture dynamics within multiple species of beetle, including *Silvacalles instabilis* A and B (Salces-Castellano et al., 2020, 2021; Noguerales et al., 2024; Noguerales & Emerson, 2025). Genetic variation within cloud forest beetle species is frequently structured into West and East populations, highlighting a dominant topoclimatic feature that has promoted cycles of population isolation and secondary contact during the Quaternary (Salces-Castellano et al., 2020). Contemporary patterns of gene flow across the area of secondary contact highlight how admixture can emerge locally (Salces-Castellano et al. 2020).

Within the broader regional genetic structure of species identified by Salces-Castellano et al. (2020), finer-scale geographical structuring of genetic variation within regions is not uncommon (Salces-Castellano et al., 2021, Noguerales et al., 2024; Noguerales & Emerson, 2025), and is indeed apparent within *Silvacalles instabilis* B, which presents strong population substructure within the very east of its range (Figure S8). Such intra-regional structure is likely driven by past or present environmental barriers to gene flow related to hydric stress landscapes (Salces-Castellano et al., 2020). With regard to the western origin and inferred isolation of the hybrid species, several species of beetle provide evidence for past barriers to dispersal that are geographically coincident with this inference. Geomitopsis franzi (Staphylinidae) presents East-West structuring consistent with that observed by Salces-Castellano et al. (2020), with further structure to the west (Perez-Delgado et al., 2022). This western structure involves three deeply divergent mtDNA lineages, with secondary contact between two of them occurring approximately 800 meters west-south-west of site MOQ (see Figure 1b). Genome-wide nuclear genetic variation with the beetle species Tarphius canariensis (Zopheridae) also reveals strong geographic structuring of genetic variation between site MOQ and further sampling 800 meters west-south-west of MOQ (Noguerales & Emerson, in prep.).

Geomitopsis franzi and Tarphius canariensis highlight a dynamic of geographic isolation and secondary contact within the western limits of the Anaga cloud forest. This is geographically coincident with where nuclear genomic data for *Silvacalles instabilis* predicts the origin of *S. instabilis* C, thus providing a mechanistic escape from the apparent paradox of Blanckaert and Bank (2018). Admixture of parental populations and a resulting hybrid swarm in the western limits of the Anaga cloud forest, with coincident or subsequent isolation from eastern populations of the parental species during glacial conditions, would provide time for reciprocal sorting of genetic incompatibilities of parental species. Inferences from both demographic modelling and individual-based simulations are consistent with hybrid origin over such a time scale. Higher cloud forest connectivity during interglacial warming (Salces-Castellano et al., 2021) would facilitate subsequent sympatry among all three species through westward range expansion of parental species and eastward range expansion of the hybrid species. It is also plausible that admixture and isolation of *S. instabilis* C occurred further west of the Anaga peninsula, either driven directly by the dramatic flank collapses and volcanic activity across the north of the island (García-Olivares et al., 2017) or indirectly through topoclimatic phenomena (Noguerales et al., 2024).

Implicit within this scenario is the local breakdown of RI between parental species *Silvacalles instabilis* A and B in the western limits of Anaga, while RI is maintained between more eastern populations. This implies geographic variation in hybridisation dynamics, something that has been observed across different hybrid zones between Hermit and Townsend’s warblers (Dendroica occidentalis and D. townsendi; Rohwer & Martin, 2007) and varieties of the common snapdragon (Antirrhinum majus subspecies majus variety pseudomajus and A. m. m. var striata; Pal et al., 2025). In the case of *S. instabilis*, a potential explanation for geographical variation of historical gene flow between parental species A and B is provided by the species pair, Tarphius canariensis and T. simplex, both codistributed with *S. instabilis*. Higher historical gene flow between western sympatric populations of the two Tarphius species, relative to eastern sympatric populations, coincides with a west-east gradient of increasing genomic divergence between populations of both species, resulting from east-west range expansions across the peninsula for both (Noguerales & Emerson, 2025). Thus, similar to both species of Tarphius, increased RI between *S. instabilis* A and B may have been facilitated by the emergence of higher genomic divergence through range expansion dynamics (Excoffier et al., 2009; Hewitt, 2004).

### Hybridisation as a facilitator of speciation?

Our data provides strong evidence for a hybrid origin for *Silvacalles instabilis* C, and for its status as a biological species. However, determining a causative role for hybridisation in the evolution of RI between *S. instabilis* C and its parental species is elusive, and highlights a more general challenge to understand the extent to which hybridisation may play a role in speciation. Hybrid speciation theory suggests that HHS times may range from tens of generations (Schumer et al., 2015) to many thousands of generations (Blanckaert & Bank, 2018), depending upon the evolutionary characteristics of genomic incompatibilities, linkage architecture and recombination rates. Accepted cases of HHS are associated with the evolution of RI over relatively short time scales (Schumer et al., 2014), where hybridisation is likely to have been a driving agent of speciation through direct selection for hybrid phenotypes (e.g., Gompert et al. 2006; Barrera-Guzmán et al., 2018; Lamichhaney et al., 2018), or background selection against genomic incompatibilities (e.g., Schumer et al., 2015). Evolution of RI over longer timescales, as might be expected when BDM incompatibilities have evolved neutrally, is suggested to be potentially more common than has been thought (Long & Rieseberg, 2024). For BDM incompatibilities that have evolved neutrally, this is theoretically possible, but requires strong isolation of hybrid populations from parental species (Blanckaert & Bank, 2018). The evolution of *S. instabilis* C can be explained within such a dynamic through a period of geographic isolation of the hybrid population from the parental species. However, given the fate of parental BDM incompatibility alleles is also nuanced by genetic architecture and population size (Blanckaert & Bank, 2018), the role of hybridisation as a driver RI between *S. instabilis* C can only be speculated upon. Shifting the focus from hybridisation as a driver to a facilitator of speciation, where hybridisation provides an initial source of partial RI that divergence can build upon, may therefore be conceptually more relevant.

Considering hybridisation as not just a driver, but also a facilitator of speciation, recognises the role of hybridisation within a spectrum, ranging from a primary driver, through a role of facilitation, to no appreciable role in the establishment of RI. While such a framework does not lessen the challenges to address and quantify the role of hybridisation in the evolution of RI, it does embrace the concerns stemming from the limitations that overly strict criteria may impose on the understanding of general principles (Nieto Feliner et al., 2017). By considering hybridisation as a facilitator of speciation, one can ask what conditions might favour the formation of hybrid populations in the first place, a necessary prerequisite to facilitation. In the case of *Silvacalles instabilis* C, it is noteworthy that both parental range sizes have been strongly impacted by Quaternary climate oscillations, and that dispersal ability is sufficiently limited to maintain genetic isolation among allopatric populations. A similar dynamic is suggested to explain the origins of a hybrid swarm between cryptic species of the torpedo scad (Megalapsis cordyla) in the western pacific (Muto et al., 2025). Indeed, it is suggested that HHS may simply reflect geographical distributions that are conducive to the formation of hybrid populations (Blanckaert & Bank, 2018), where the Italian sparrow (Passer italiae; Runemark et al., 2018), the South American fur seal (Arctocephalus australis; Lopes et al., 2023) and the Aigoual grasshopper (Chorthippus saulcyi algoaldensis; Noguerales & Ortego, 2022), also prove compelling examples. If speciation is facilitated by the formation of hybrid populations, then time to speciation (i.e., RI=1) for such populations from their parental species should be less than that of the parental species through cladogenesis. Our coalescent simulations are consistent with this. Despite hybridisation between parental species being approximately four times more recent than cladogenetic speciation between parental species, RI of the hybrid species already approaches that observed between parental species. While we are unable to say if hybridisation drove speciation within *S. instabilis*, it would seem likely that it has played an important role.

## Conclusions

*Silvacalles instabilis* provides the most direct and compelling evidence to date for reproductive isolation between a species of hybrid origin and both parental species. The parental species *S. instabilis* A and *S. instabilis* B are estimated to have diverged approximately 800,000 years ago, with subsequent hybridisation giving rise to *S. instabilis* C approximately 200,000 years ago. An ancient rather than recent hybrid origin for *S. instabilis* C is also supported by genome-wide nuclear estimates of heterozygosity and individual-based simulations of hybrid speciation. Based on geographic patterns of genomic relatedness between hybrid and parental individuals, the most plausible explanation is that *S. instabilis* C evolved within a hybrid swarm that remained geographically isolated from both parental species long enough for reproductive isolation to emerge. The phenotypic, ecological and temporal dimensions of speciation within *S. instabilis* suggest the need for a more nuanced consideration of the role of hybridisation in speciation. Much focus has been placed on hybridisation as a driving agent of speciation. We argue that hybridisation may also plays a facilitative role. This recognises the speciation relevance of hybridisation within a spectrum, ranging from a primary driver, through a gradient of facilitation, to the other extreme of no appreciable role in the establishment of RI. We suggest that isolated hybrid swarm populations are likely to evolve RI faster than allopatric isolates of parental populations, and that this may be a useful framework to understand hybridisation as a facilitator of speciation.

## AUTHOR CONTRIBUTIONS

BCE and VN conceived the original idea, designed the research and led the study. VN led the analyses, with contributions from ME, SLG and SS. BCE and VN wrote the manuscript. All authors contributed critically to the draft and gave final approval for publication.

## ACKNOWLEDGEMENTS

We wish to thank the ‘Centro de Supercomputación de Galicia (CESGA)’ and the Teide High-Performance Computing facility (teideHPC) provided by the ‘Instituto Tecnológico y de Energías Renovables (ITER), S.A.’ for access to computer resources. We thank Antonio J. Pérez-Delgado for ddRADseq library preparation. We also extend our gratitude to Heriberto López, Eduardo Jiménez, Daniel Suárez, Antonio Machado and Antonio Pérez-Delgado for assistance with fieldwork, specimen sorting, taxonomic identification and genomic library preparation. We thank Carmelo Andújar for fieldwork, specimen sorting, DNA extraction and mtDNA sequencing. Fieldwork was supported by the ‘Cabildo de Tenerife’ (Expte: AFF17/23, N° Sigma: 2023-00133). This work was supported by projects CGL2017-85718-P and PID2020-116788GB-I00, financed by MCIN/AEI/10.13039/501100011033 and co-financed by FEDER, awarded to BC. VN was also supported by a ‘Viera y Clavijo’ postdoctoral fellowship funded by the ‘Agencia Canaria de Investigación, Innovación y Sociedad de la Información’ and the ‘Universidad de La Laguna’.

## SUPPLEMENTARY INFORMATION

### SUPPLEMENTARY METHODS

#### Methods S1. Sample collection

Specimens from the *Silvacalles instabilis* (Wollaston, 1864) species complex were collected within the cloud forest of the Anaga peninsula, located on the Canary island of Tenerife (Figure 1). To augment the ten sampling sites included originally in Salces-Castellano et al. (2020), additionally sampling was conducted giving rise to a total of 110 specimens from a total of 21 sampling sites (Figure 1; Table S1), including 106 individuals from *S. instabilis*, and 4 individuals from the sister species *S. nubilosus*, which were used as an outgroup. The geographic distance between sampling sites ranged from 0.2 km to 14 km along the dorsal ridge of the Anaga peninsula, with a maximum elevational difference between sites of 200 m (Table S1). Sampling was performed following the protocol described in Salces-Castellano et al. (2020). Specimens were preserved in 100% ethanol and stored at -20°C till DNA extraction. Sampling was undertaken with permits issued by ‘Cabildo de Tenerife’ (Expte: AFF17/23, N° Sigma: 2023-00133).

#### Methods S2. Genomic data filtering and sequence assembly

First, the overall quality of raw Illumina reads was visually inspected using fastqc version 0.11.7 (Andrews, 2010). Raw reads were then demultiplexed, quality filtered and de novo assembled using ipyrad version 0.9.96 (Eaton & Overcast, 2020). A stricter filter to remove Illumina adapter contamination (filter_adapters) was applied, and only reads with unambiguous barcodes (max_barcode_mismatch) were retained. After trimming restriction overhangs for enzymes EcoR1 and MseI (restriction_overhang), base calls with a Phred score <20 were converted into ambiguous sites (Ns) and reads with >5 Ns (max_low_qual_bases) were discarded. Afterwards, the retained reads within and across samples were clustered considering a threshold of sequence similarity of 85% (clust_threshold), and those clusters with a minimum coverage depth of less than 5 (mindepth_majrule) and a maximum coverage depth of more than 10,000 (maxdepth) were discarded. Statistical base calling was performed at a minimum depth of 6 (mindepth_statistical). Resulting loci shorter than 35 base pairs (bp) (filter_min_trim_len) and with more than 20% polymorphic sites (max_SNPs_locus) were discarded. The maximum proportion of shared heterozygous sites in a locus (max_shared_Hs_locus) was set to 50%. In a final filtering step, only loci that were present in at least 80% of the samples (min_samples_locus) were retained. This yielded a total of 5040 unlinked SNPs when including all 106 specimens, 5250 unlinked SNPS when including only the 45 individuals from lineage A, 7927 unlinked SNPs when including only the 48 individuals from lineage B, and 7022 unlinked SNPs when including only individuals of hybrid origin (13 specimens from the admixed lineage C). On average, missing data in each SNP matrix was approximately 10.88%-13.64%. According to ipyrad, estimates of sequencing error rates and heterozygosity across the 106 individuals were on average 0.00103 (SD=0.00033) and 0.01785 (SD=0.00325), respectively.

#### Methods S3. Genomic clustering analyses in structure

Nuclear relatedness among individuals and the potential signatures of admixture between them were examined using the Bayesian Markov Chain Monte Carlo (MCMC) clustering method implemented in structure version 2.3.3 (Pritchard et al., 2000). structure was run with 200,000 MCMC cycles after a burn-in step of 100,000 iterations, assuming correlated allele frequencies and admixture (Pritchard et al., 2000) and performing 10 independent runs for each value of K ancestral populations (from *K*=1 to *K*=5). The most likely number of ancestral populations was estimated after retaining the 10 runs per each K-value with the highest likelihood estimates. Convergence across runs was assessed by checking that the 10 retained replicates per K-value provided a similar solution in terms of individual probabilities of assignment to a given ancestral population (q-values; Gilbert et al., 2012). As recommended by Gilbert et al. (2012) and Janes et al. (2017), two statistics were used to interpret the number range of ancestral populations (K) that best describes our data: log probabilities of Pr(X|*K*) (Pritchard et al., 2000) and Δ*K* (Evanno et al., 2005), both calculated in structure harvester (Earl & vonHoldt, 2012). Finally, the Greedy algorithm in clumpp version 1.1.2 was used to align replicated runs of structure for the same K-value (Jakobsson & Rosenberg, 2007).

#### Methods S4. Phylogenomic inferences

Phylogenetic relationships among all individuals were reconstructed using a maximum-likelihood (ML) approach as implemented in raxml version 8.2.12 (Stamatakis, 2014). This analysis was performed on a matrix of concatenated SNPs (i.e., including all SNPs per RAD locus), which was edited with the r package phrynomics (B. Banbury, http://github.com/bbanbury/phrynomics) to remove invariant and non-binary SNPs, resulting in a dataset of 20,407 concatenated SNPs. The ascertainment bias correction based on the conditional likelihood method (Lewis, 2001) and the GTR-GAMMA model of nucleotide evolution were applied. The best-scoring maximum likelihood tree and node supports were estimated through 100 rapid bootstrap replicates. The 4 individuals from the sister species *Silvacalles nubilosus* (Table S1) were used as an outgroup.

Phylogenetic relationships among higher-order genetic groups were reconstructed using snapp version 1.5.2 (Bryant et al., 2012), a coalescent-based method for species tree estimation, as implemented in beast version 2.6.7 (Bouckaert et al., 2014). Individuals were assigned to each of the three genetic groups (either lineage A, B or C), according to the clustering scheme observed in the global PCA and structure inferences, assuming *K*=2 (Figure 1, Figure S1). To reduce the high computational burden of snapp, a total of 20 individuals for each of the two main genetic groups (lineages A and B) were selected, encompassing the overall within-lineage genetic variation observed in their respective clustering analyses (Figures S7-S8). Regarding lineage C, the individual classified as an *F*_1_ hybrid with lineage B was excluded (Figure 1ac), yielding a total of 12 individuals of admixed co-ancestry. The final dataset contained 52 individuals distributed across the three demes. Phylogenetic relationships using snapp were also reconstructed by taking into account the geographical structuring of genetic variation within each of the two main lineages (lineages A and B). For each lineage, individuals were assigned to geographically coherent populations according to the clustering schemes observed in the within-lineage PCA and structure analyses, assuming *K*=2 (Figures S7-S8). Accordingly, a total of 12 individuals of high single ancestry (q-value >0.95) were selected for each of the genetic groups, corresponding to western and eastern lineage A, and western and eastern lineage B. Regarding lineage C, the aforementioned subset of 12 individuals was included. The final dataset contained 60 individuals distributed across the five demes.

For each of the two datasets, the respective .usnps file from ipyrad was edited with the r package phrynomics to remove non-binary SNPs, which resulted in matrices including 5615 and 5199 bi-allelic unlinked SNPs shared across demes, respectively. The default values for the shape (α) and inverse scale (β) parameters of the gamma prior distribution for the population size parameter (θ) were set, and the forward (u) and reverse (v) mutation rates were left to be calculated by snapp. The log-likelihood correction was applied, and the coalescent rate was set to be sampled, with the remaining parameters left as default. Two independent runs using different starting seeds for 1 million MCMC generations were performed, sampling every 1000 steps (∼1000 genealogies). The log files were examined in tracer version 1.7 (Rambaut et al., 2018) to check stationarity and convergence of the chains and confirm that effective sample sizes (ESS) for all parameters were >200. The software logcombiner version 2.4.7 (Drummond & Rambaut, 2007) was used to remove 10% of trees as burn-in and combine tree and log files for replicated runs. Finally, treeannotator version 2.4.7 (Drummond & Rambaut, 2007) and densitree version 2.2.6 (Bouckaert, 2010) were used to obtain the maximum credibility trees and displayed the full set of retained trees, respectively.

#### Methods S5. Introgression analyses using ABBA-BABA tests

Potential signatures of gene flow between non-sister taxa were examined using four-taxon ABBA-BABA tests based on the D-statistics (Durand et al., 2011), as implemented in ipyrad (Eaton & Overcast, 2020). Standard deviations of D-statistics were obtained through 1000 bootstrap replicates. Assuming the topology retrieved in raxml (Figures S1-S2), firstly all individuals were assigned to one of the three lineages A, B and C, according to the clustering pattern observed in the global PCA and structure analyses. Then, ABBA-BABA tests were run using the same subset of 52 individuals used in snapp for reconstructing the phylogenetic relations among the three higher-order genetic groups (Figures S2a). Finally, ABBA-BABA tests were run using the same subset of 60 individuals used in snapp to take into account the geographical structuring of genetic variation within each of the two main lineages (lineages A and B) (Figures S2b). For all analyses, the 4 individuals of *Silvacalles nubilosus* were used as an outgroup.

#### Methods S6. Testing alternative models of divergence and hybridisation in fastsimcoal

To evaluate the relative statistical support for each of the alternative demographic scenarios (Figure S3, Tables S3), the composite likelihood of the observed data given a specified model was estimated in fastsimcoal version 2.5.2.21 (Excoffier et al., 2021) using the site frequency spectrum (SFS). These analyses were conducted using the same subset of 52 individuals selected for reconstructing phylogenetic relationships in snapp among the higher-order genetic groups. The folded joint SFS was calculated using easysfs version 0.0.1 (I. Overcast, https://github.com/isaacovercast/easySFS), considering a single SNP per locus to avoid the effects of linkage disequilibrium. Each of the three genetic groups was downsampled to 180% of individuals to remove all missing data for the calculation of the SFS, minimise errors with allele frequency estimates, and maximise the number of variable SNPs retained. The final SFS contained 4272 variable SNPs. Because invariable sites were not included in the SFS, the ‘removeZeroSFS’ option in fastsimcoal was applied. Accordingly, the effective population size (*N*_e_) for one of the demes (lineage A) was fixed to enable the estimation of other parameters (Papadopoulou & Knowles, 2015; Excoffier et al., 2021; Noguerales & Ortego, 2022). To this end, the parameter *N*_e_ was calculated from observed estimates of nucleotide diversity (π) and mutation rate per site per generation (μ), where *N*_e_=π/4μ. The software dnasp version 6.12.03 (Rozas et al., 2017) was used to estimate π for the fixed deme using phased data from polymorphic and non-polymorphic loci contained in the .allele file from ipyrad. A mutation rate per site per generation of 2.8×10 (Keightley et al., 2014), as estimated for the butterfly Heliconius melpomene, was considered.

A total of 100 independent replicated runs were conducted for each model, considering 100,000-250,000 simulations for the calculation of the composite likelihood, 10-40 expectation-conditional maximisation (ECM) cycles, and a stopping criterion of 0.001 (Excoffier et al., 2021). An information-theoretic model selection approach based on Akaike’s information criterion (AIC) was applied to determine the probability of each model given the observed data (Burnham & Anderson, 2002; e.g., Thomé & Carstens, 2016). The replicated run for each model with the highest composite likelihood was selected to calculate its respective AIC score, as detailed in Thomé and Carstens (2016). Then, AIC values were rescaled across models, calculating the difference between the AIC value of a given model and the minimum AIC obtained among all competing models (i.e., the best model has ΔAIC=0). The point estimates of the different demographic parameters for the most-supported model were selected from the run with the highest maximum composite likelihood. Finally, the confidence intervals of the parameter estimates (based on the percentile method; e.g., de Manuel et al., 2016) were estimated under the most-supported model through 100 parametric bootstrap replicates by simulating SFS from the maximum composite likelihood estimates and re-estimating parameters each time (Excoffier et al., 2021).

#### Methods S7. Individual-based simulations of hybrid speciation

For simulations with linkage, genetic markers were randomly arrayed onto n chromosomes, each with dimensions 0,1. To simulate free recombination, each marker was assigned to separate chromosomes. For the scenarios with 20 chromosomes and 1 chromosome, each chromosome pair experienced 1 crossover per generation. For the tight linkage scenario, the per-chromosome recombination rate was reduced to 0.05, and for the scenario with no recombination no crossovers were allowed to occur. To allow for low rates or migration, parental individuals were introduced in the population as rate m. Because the population size is relatively small, low migration was simulated by only introducing migrants at specific intervals.

For example, to achieve an overall rate of *m*=0.001, a parental migrant was introduced every 5 generations.

## SUPPLEMENTARY TABLES

**Table S1.**
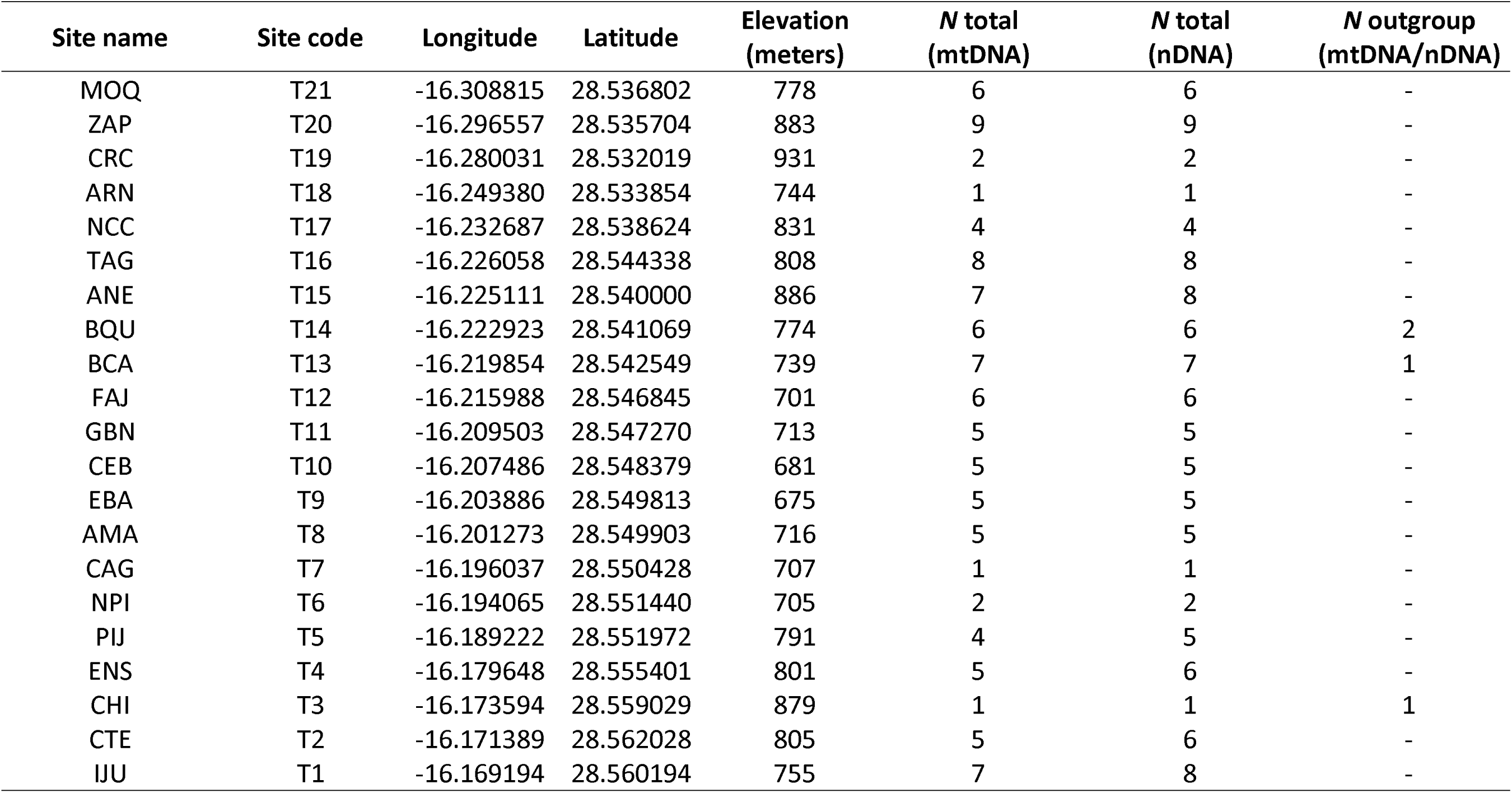
Geographic coordinates and elevation for each of the sampling sites. For each site, the total number of individuals from the *Silvacalles instabilis* species complex with mitochondrial (mtDNA) and genome-wide nuclear (nDNA) data is indicated. Four individuals belonging to the sister species *Silvacalles nubilosus* were used as an outgroup, for which mtDNA and nDNA data was also generated.

**Table S2.** Analyses of introgression using *D*-statistics (ABBA-BABA tests), assuming the topological inferences obtained in raxml and snapp. The value of the *D*-statistics, its standard deviation (SD), *Z*-score and *P*-value are provided for each test. Assuming that the sister taxa P_1_ and P_2_ diverged from P_3_, the *D*-statistic is used to test the null hypothesis of no introgression (*D*-statistics=0) between P_3_ and P_1_ or P_2_. *D*-statistics significantly different from 0 indicate gene flow between P_2_ and P_3_ (*D*-statistics > 0) or between P_1_ and P_3_ (*D*-statistics < 0). Analyses were performed (i) using all individuals in each higher-order genetic group, corresponding to lineages A, B and C (tests 1-2), (ii) including the same subset of 20 individuals from each lineage A and B (tests 3-4), as used for snapp and fastsimcoal (see Supplementary Methods S4-S6), and (iii) taking into account the structuring of genetic variation within the higher-order genetic groups, and therefore assigning individuals into geographic populations of single co-ancestry, according to the clustering schemes observed in the within-lineage analyses (tests 5-12). All tests were repeated after excluding the *F*_1_individual between lineages B and C (Figure 1, Figure 2), giving rise to a P_2_ group composed of 12 individuals. For all analyses, the four individuals from *S. nubilosus* were used as an outgroup.

| Test | $P_1$ | $P_2$<br>(hybrid) | $P_3$ | *D*-statistics | SD | Z-score | *P*-value |
| --- | --- | --- | --- | --- | --- | --- | --- |
| 1 | Lineage B (all ind.) | Lineage C (12 ind.) | Lineage A (all ind.) | 0.084 | 0.017 | 5.121 | <0.001 |
| 2 | Lineage B (all ind.) | Lineage C (all ind.) | Lineage A (all ind.) | 0.082 | 0.016 | 5.563 | <0.001 |
| 3 | Lineage B (20 ind.) | Lineage C (12 ind.) | Lineage A (20 ind.) | 0.088 | 0.017 | 5.162 | <0.001 |
| 4 | Lineage B (20 ind.) | Lineage C (all ind.) | Lineage A (20 ind.) | 0.085 | 0.017 | 5.063 | <0.001 |
| 5 | Western lineage B (12 ind.) | Lineage C (12 ind.) | Western lineage A (12 ind.) | 0.073 | 0.018 | 4.088 | <0.001 |
| 6 | Eastern lineage B (12 ind.) | Lineage C (12 ind.) | Western lineage A (12 ind.) | 0.112 | 0.018 | 6.388 | <0.001 |
| 7 | Western lineage B (12 ind.) | Lineage C (all ind.) | Western lineage A (12 ind.) | 0.070 | 0.017 | 4.115 | <0.001 |
| 8 | Eastern lineage B (12 ind.) | Lineage C (all ind.) | Western lineage A (12 ind.) | 0.110 | 0.017 | 6.370 | <0.001 |
| 9 | Western lineage B (12 ind.) | Lineage C (12 ind.) | Eastern lineage A (12 ind.) | 0.110 | 0.018 | 6.118 | <0.001 |
| 10 | Eastern lineage B (12 ind.) | Lineage C (12 ind.) | Eastern lineage A (12 ind.) | 0.074 | 0.019 | 3.966 | <0.001 |
| 11 | Western lineage B (12 ind.) | Lineage C (all ind.) | Eastern lineage A (12 ind.) | 0.108 | 0.017 | 6.222 | <0.001 |
| 12 | Eastern lineage B (12 ind.) | Lineage C (all ind.) | Eastern lineage A (12 ind.) | 0.072 | 0.018 | 3.879 | <0.001 |

**Table S3.**
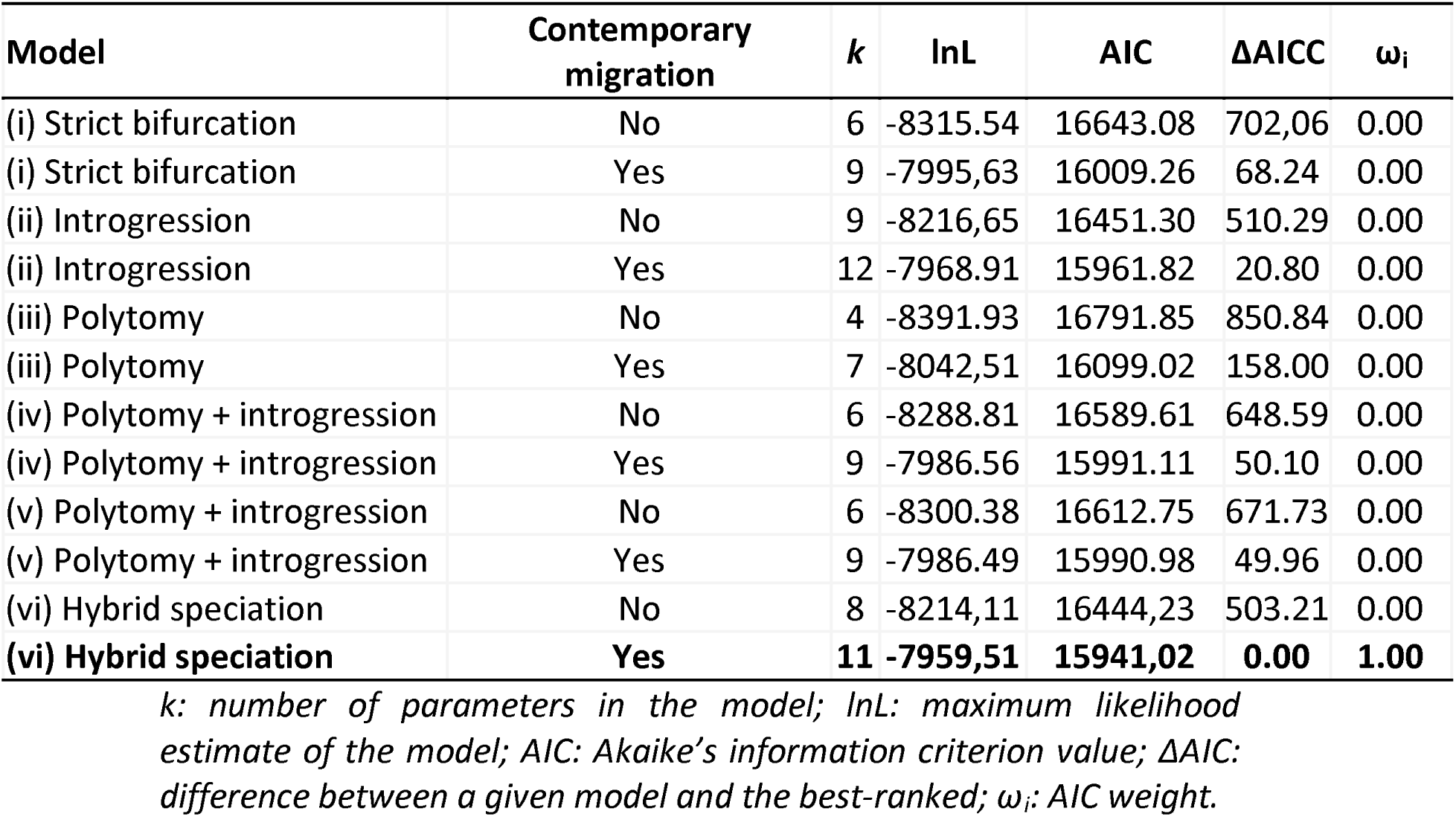
Comparison of the alternative demographic scenarios of divergence and hybridisation within the *Silvacalles instabilis* species complex, tested using fastsimcoal (Figure S3). The most-supported model is highlighted in bold (Figure 3). Each topological model was evaluated either assuming contemporary symmetrical migration among all demes or the lack of it, yielding a total of twelve competing demographic scenarios.

## SUPPLEMENTARY FIGURES

**Figure S1.**
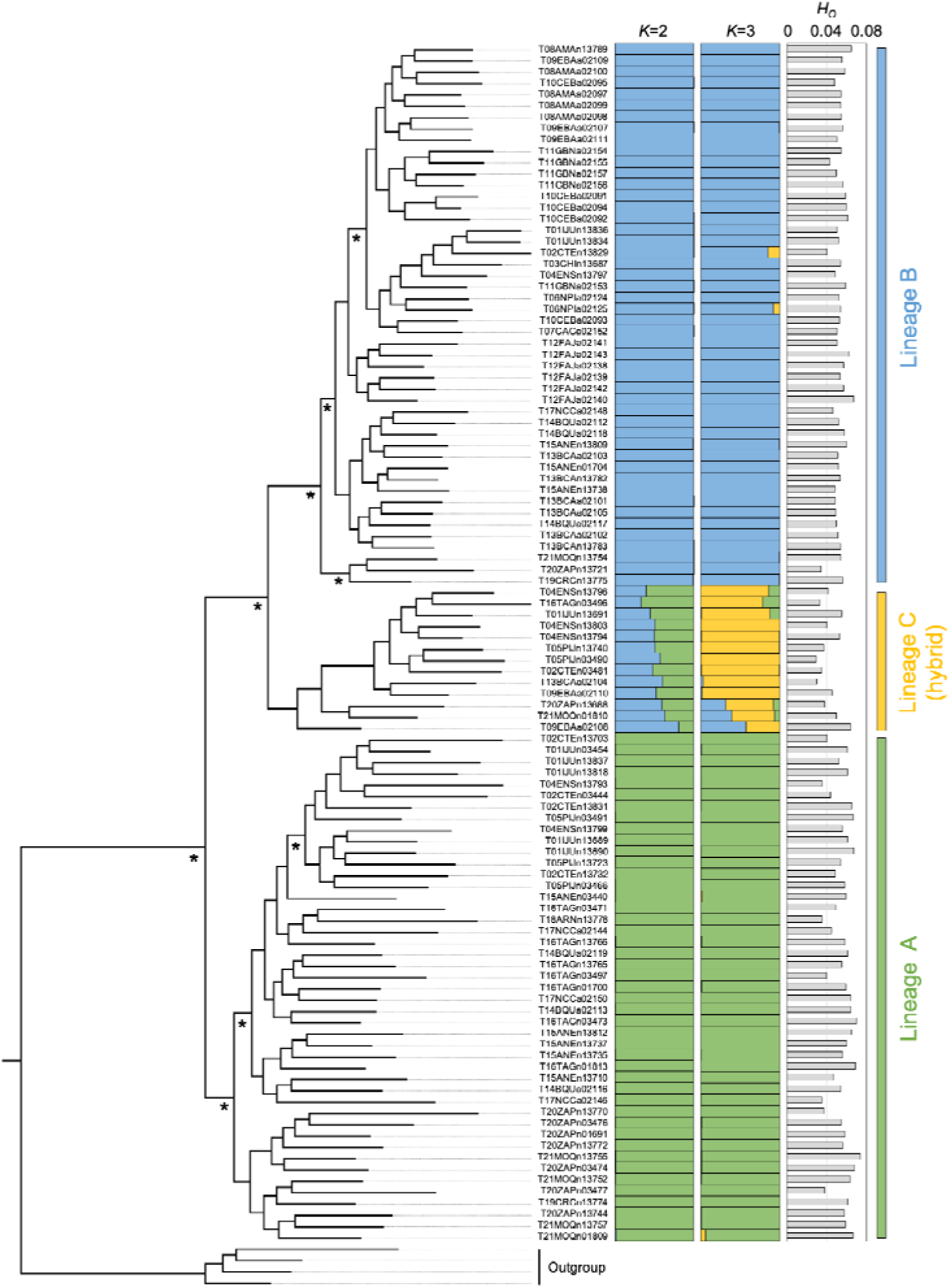
Phylogenetic relationships among individuals of the *Silvacalles instabilis* specie complex, as inferred from genome-wide nuclear data using raxml. Asterisks on the tree denote fully supported nodes (bootstrap value >99). Colour barplots represent the individual-based ancestry coefficients as inferred in structure, assuming two and three ancestral population (*K*=2-3). For each individual, the right grey bar represents its genetic variability as measured using the observed heterozygosity (*H*_O_) parameter. Four individuals belonging to the sister species *S. nubilosus* were used as an outgroup. Sites codes are as in Table S1, with colours as in Figure 1.

**Figure S2.**
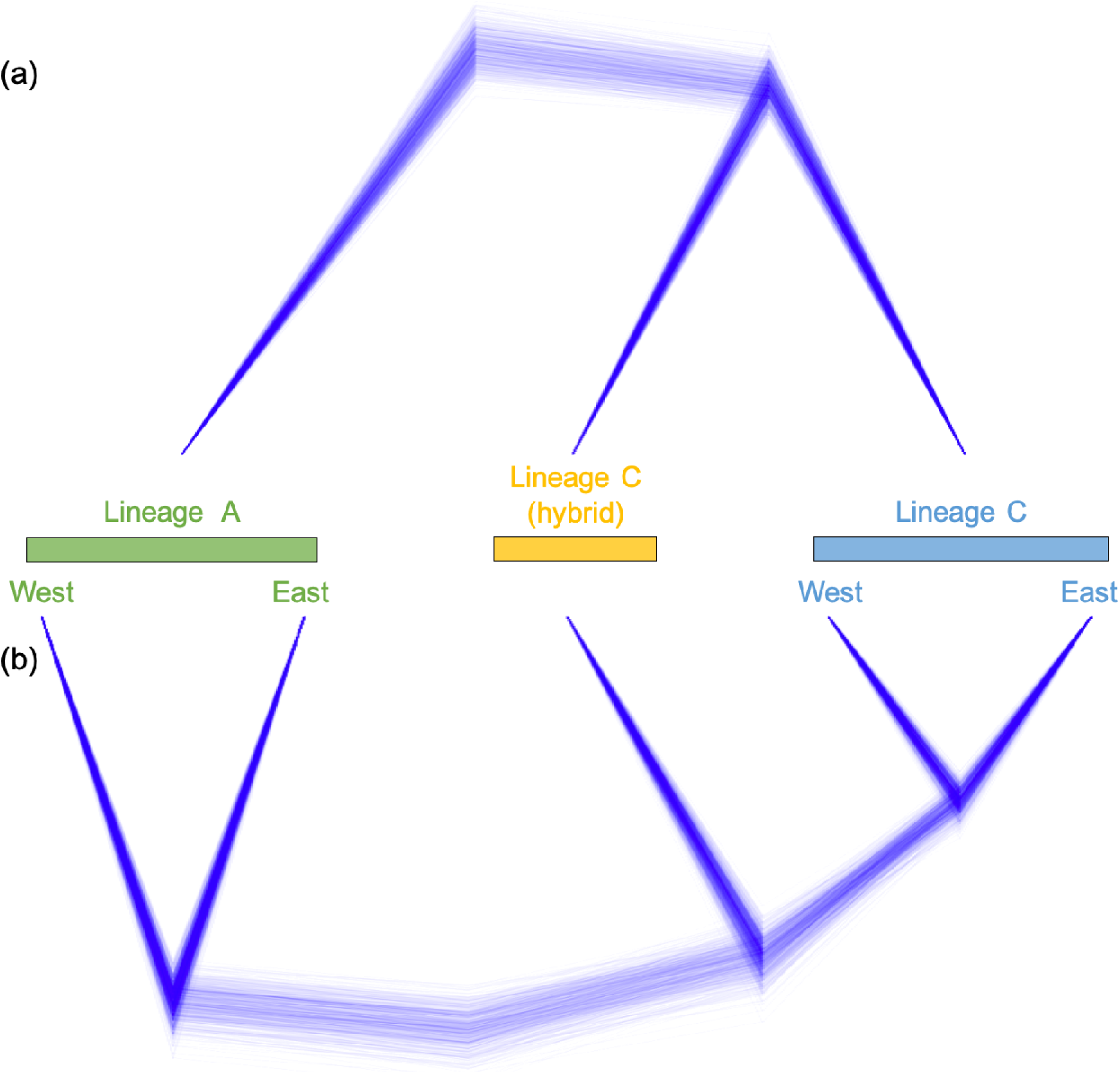
Phylogenetic relationships as inferred in snapp among (a) the three higher-order genetic groups within the *Silvacalles instabilis* species complex, and (b) their geographic populations according to the clustering schemes observed in the within-lineage analyses by structure. All nodes are fully supported.

**Figure S3.**
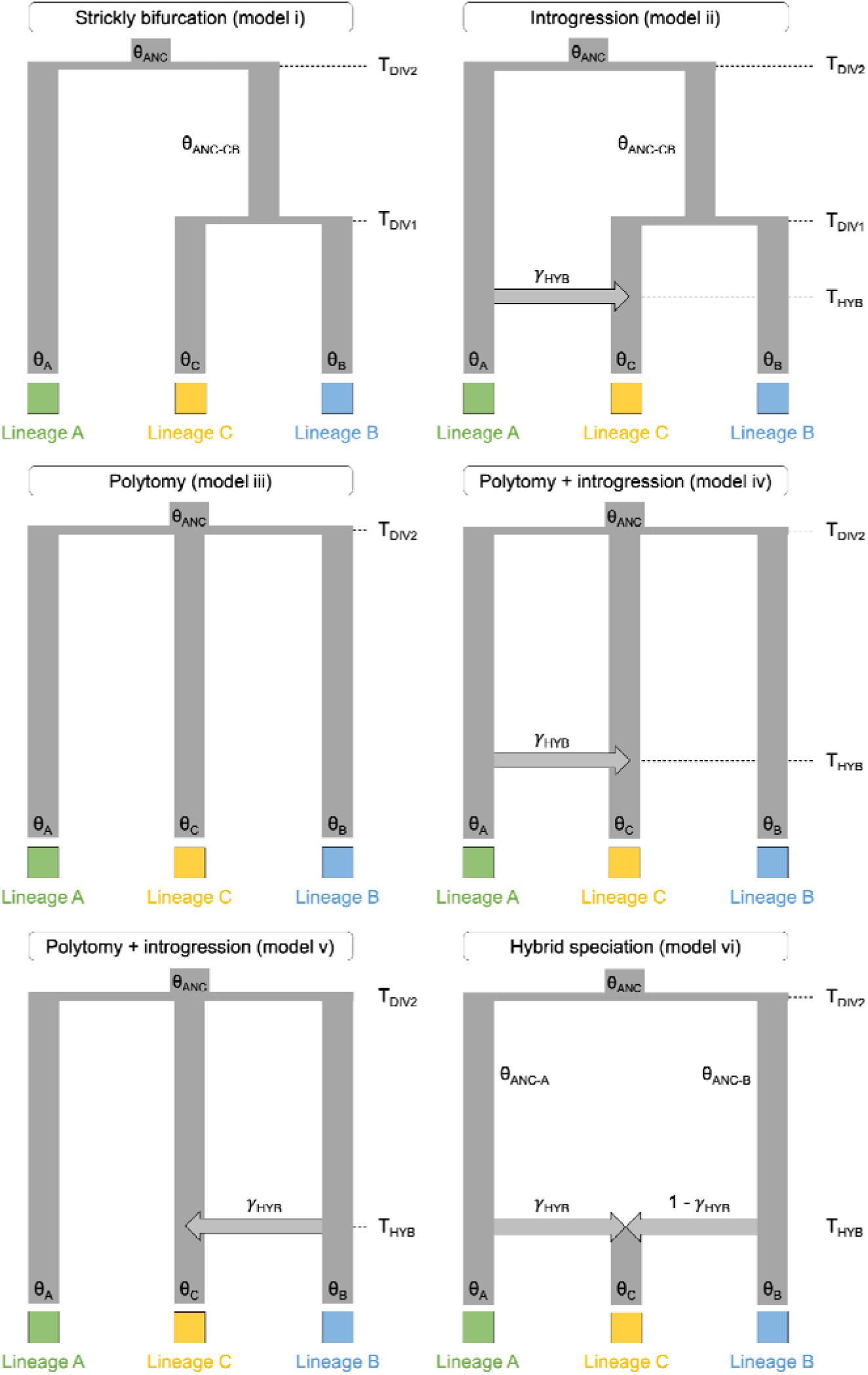
Alternative demographic scenarios of divergence and hybridisation within the *Silvacalles instabilis* species complex, tested using fastsimcoal. Model parameters include ancestral (θ_ANC_, θ_ANC-A_, θ_ANC-B_, and θ_ANC-CB_) and contemporary (θ_A_, θ_B_, and θ_C_) effective population sizes, timing of divergence (T_DIV1_, T_DIV2_), timing of hybridisation (T_HYB_), and contribution coefficient (γ_HYB_). These six models were also evaluated either assuming symmetrical contemporary migration (*m*_I-J_) among all demes (not shown on hypothetical scenarios).

**Figure S4.**
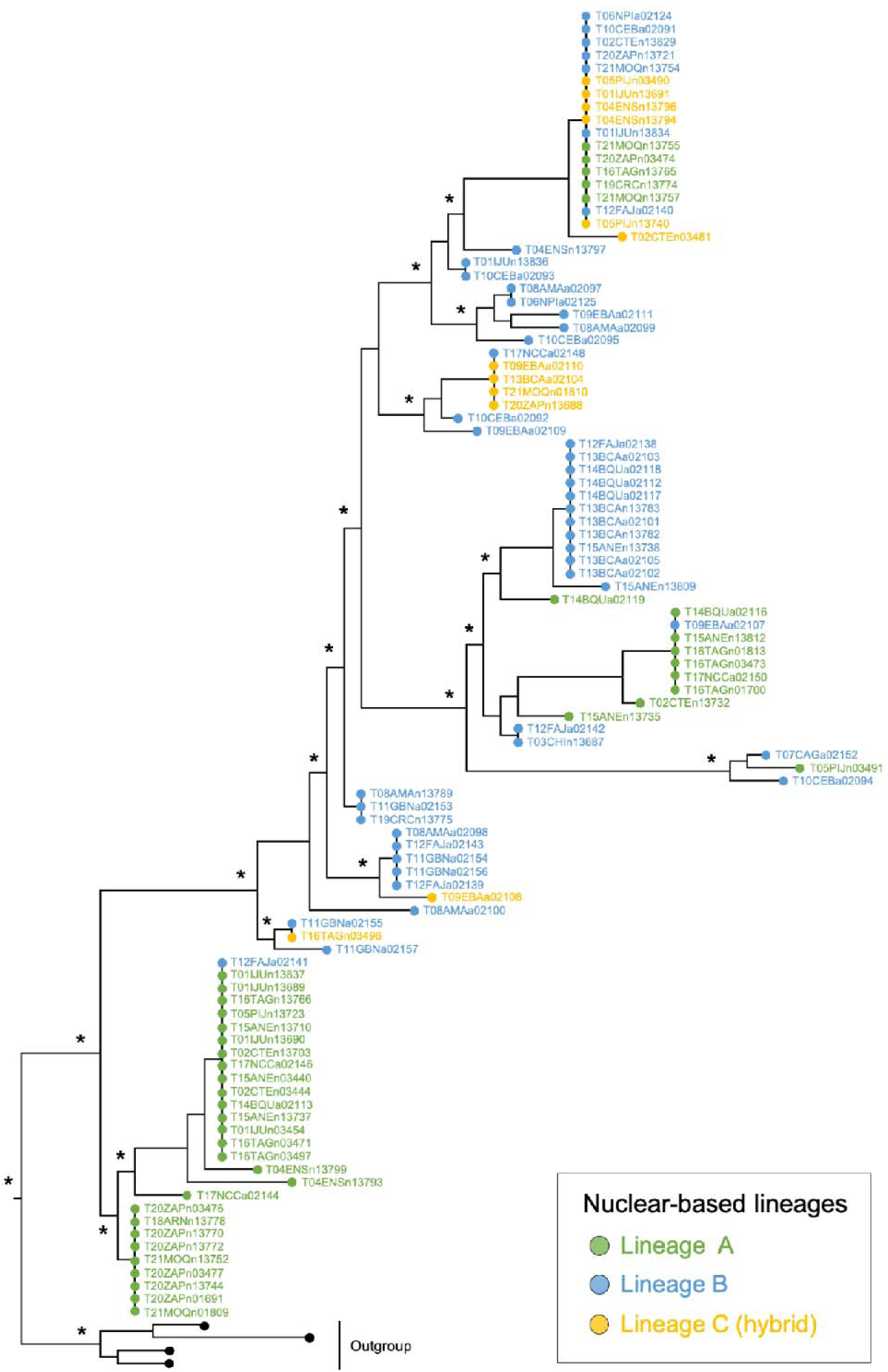
Phylogenetic relationships among individuals of the *Silvacalles instabilis* specie complex, as inferred from the barcode region of the mitochondrial DNA (mtDNA) cytochrome c oxidase subunit I (COI) using iq-tree. Associations between mtDNA haplotypes and higher-genetic groups based on genome-wide nuclear data are mapped onto the mtDNA tree. Asterisks on the tree denote well-supported nodes (bootstrap value >85). Four individuals belonging to the sister species *S. nubilosus* were used as an outgroup. Colours as in Figure 1.

**Figure S5.**
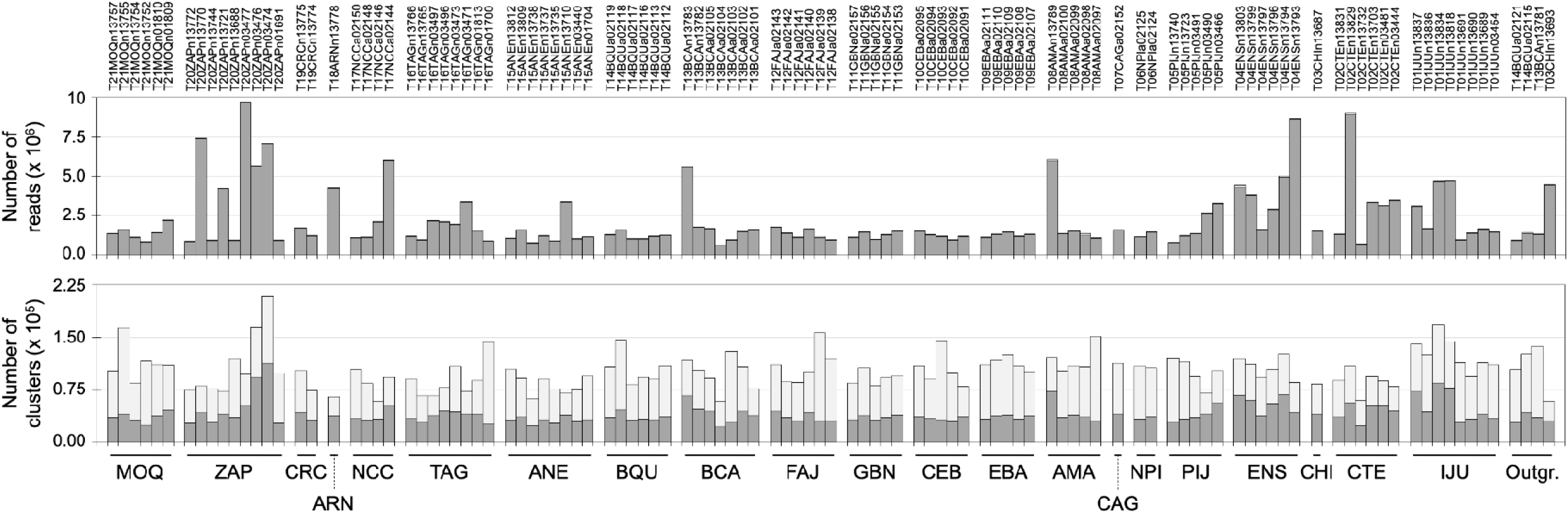
The upper plot shows the number of reads per individual after the different quality filtering steps conducted in ipyrad. The cumulative stacked bars in the bottom plot summarise the number of sequence clusters before and after the assembly and filtering steps in ipyrad. Within each bar, the pale grey colour represents the clusters that were discarded during the filtering procedure, with the dark grey colour representing the total number of retained clusters. Site codes as in Table S1.

**Figure S6.**
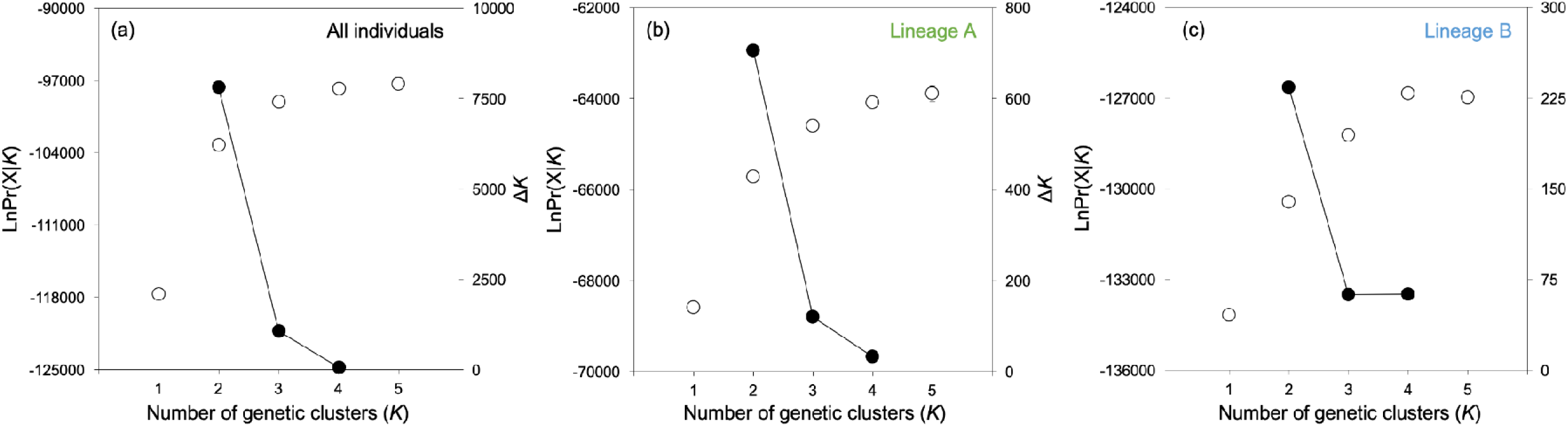
Mean (±SD) log probability of the data (LnPr(X|*K*); (left axes, open dots and error bars) and the magnitude of Δ*K* (right axes, black dots and continuous line) for each value of *K*-value estimated over 10 structure runs for three different analyses including (a) all individuals from the *Silvacalles instabilis* species complex, (b) only individuals assigned to lineage A, and (c) only individuals assigned to lineage B (Figure 1).

**Figure S7.**
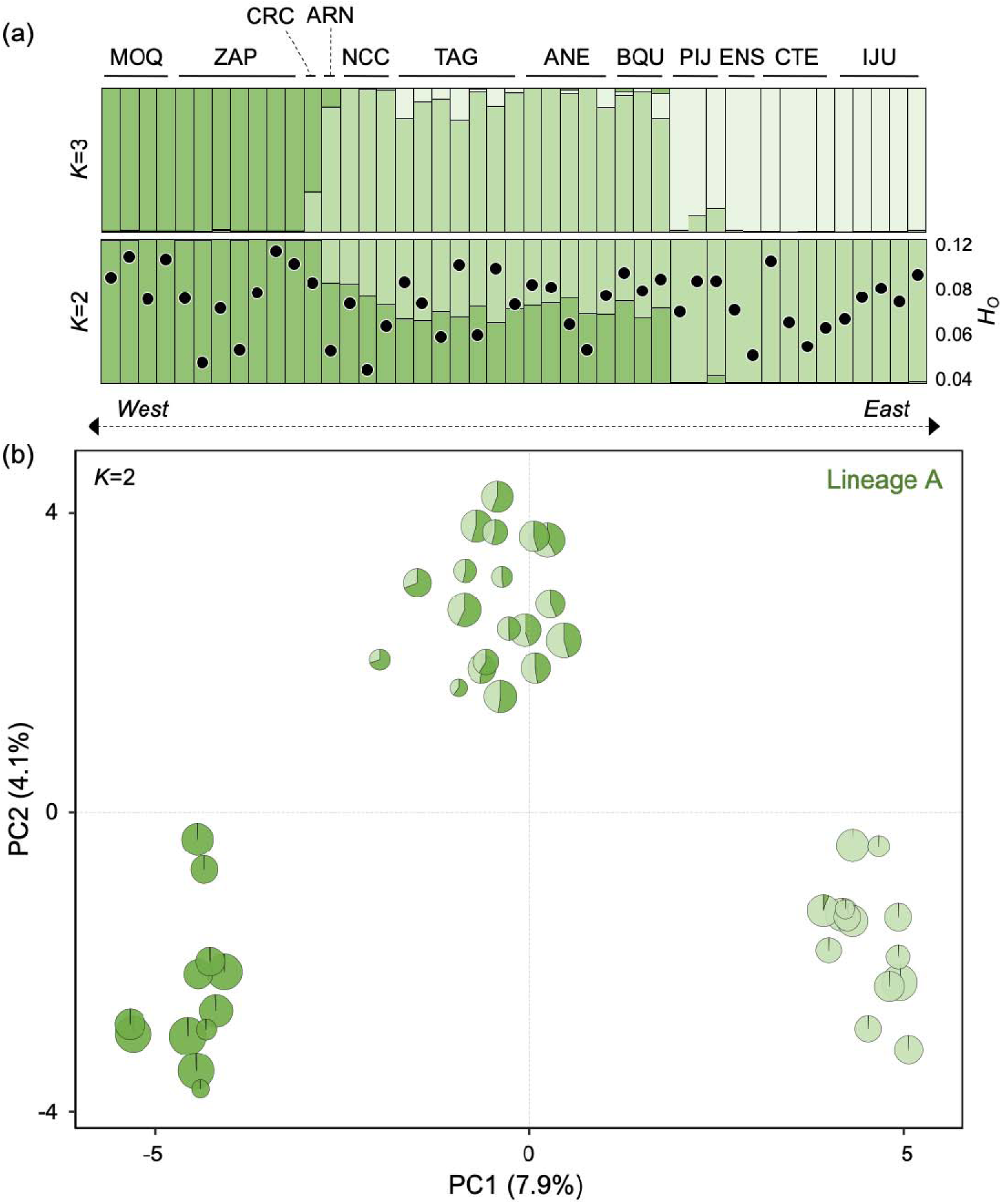
Geographical structuring of nuclear genomic variation within lineage A, as inferred with structure (panel a) and a principal component analysis (PCA) (panel b). In panel (a), structure bar plots depict the ancestry coefficients per individual assuming two and three ancestral populations (*K*). Thin vertical black lines separate individuals, which are partitioned into *K*-coloured segments representing the probability of belonging to a given ancestral population. Within the *K*=2 bar plot, black circles represent the level of genetic variability pe individual as measured using the observed heterozygosity (*H*_O_) parameter. Site codes as in Table S1. In panel (b), pie charts represent the position of each individual along the two first principal components (PCs) and their respective ancestry coefficients based on structur results, assuming *K*=2. Size of pie charts represents individual-based *H*_O_.

**Figure S8.**
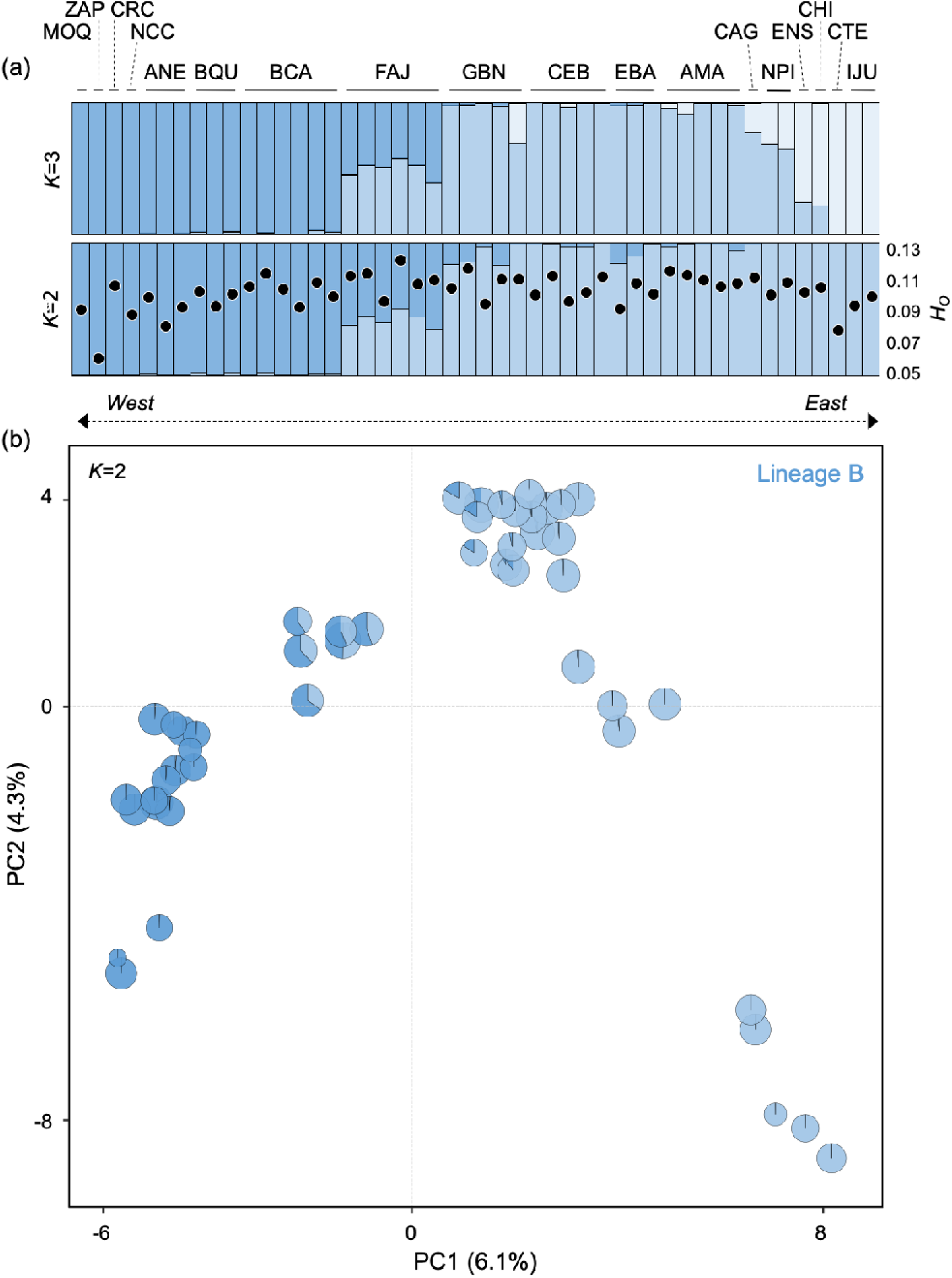
Geographical structuring of the genomic variation within lineage B, as inferred with structure (panel a) and a principal component analysis (PCA) (panel b). In panel (a), the structure bar plots depict the ancestry coefficients per individual assuming two and three ancestral populations (*K*). Thin vertical black lines separate individuals, which are partitioned into *K*-coloured segments representing the probability of belonging to a given ancestral population. Within the *K*=2 bar plot, black circles represent the level of genetic variability pe individual as measured using the observed heterozygosity (*H*_O_) parameter. Site codes as in Table S1. In panel (b), pie charts represent the position of each individual along the two first principal components (PCs) and their respective ancestry coefficients based on structur results, assuming *K*=2. Size of pie charts represents individual-based *H*_O_.

**Figure S9.**
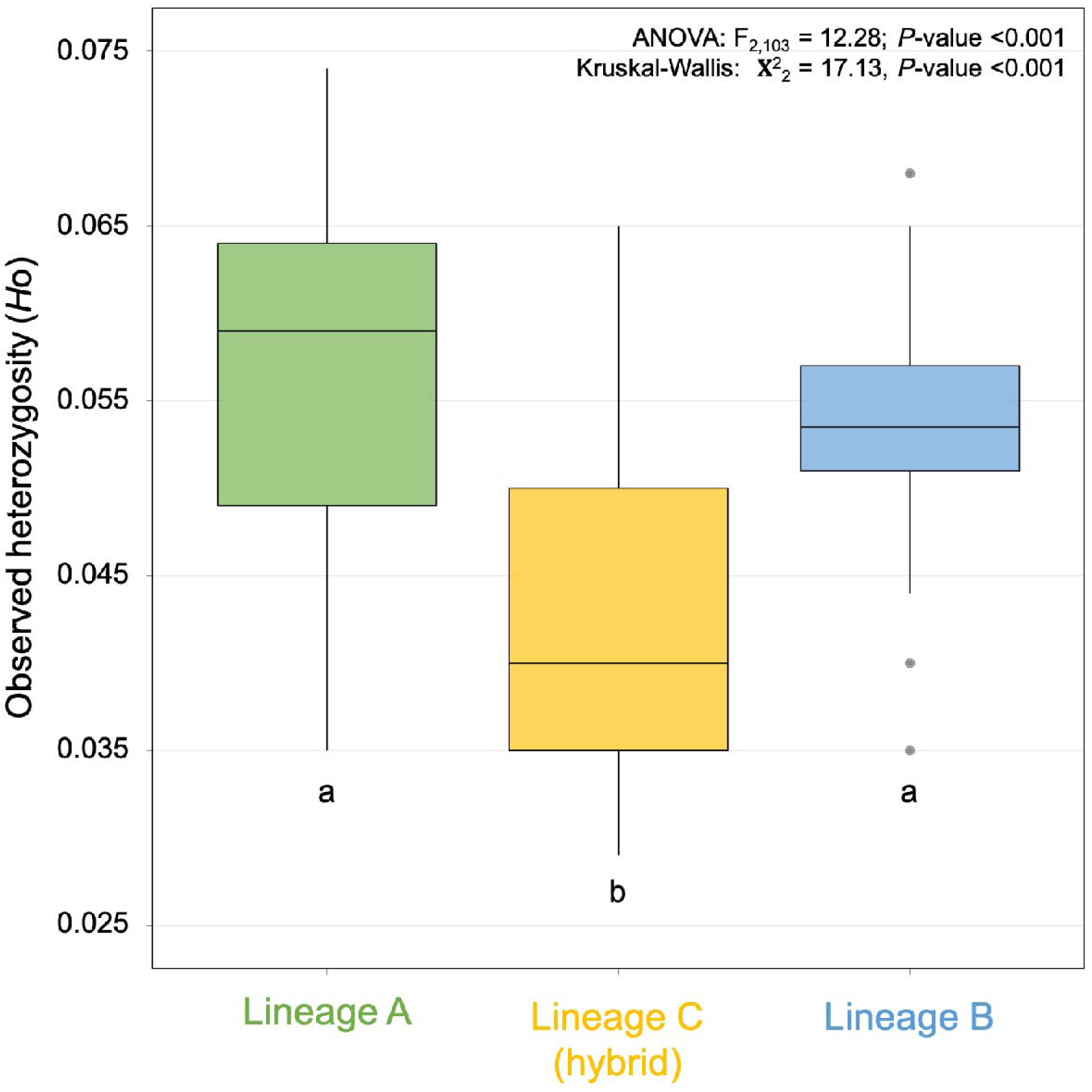
Observed heterozygosity (*H*_O_) across the higher-order genetic groups according to structure results, assuming three ancestral populations (*K*=3; Figure S1). Letters below box plot indicate that differences between the respective groups are not statistically significant (*P*-value >0.05) after post hoc tests using both the Tukey and Wilcoxon methods.

**Figure S10.**
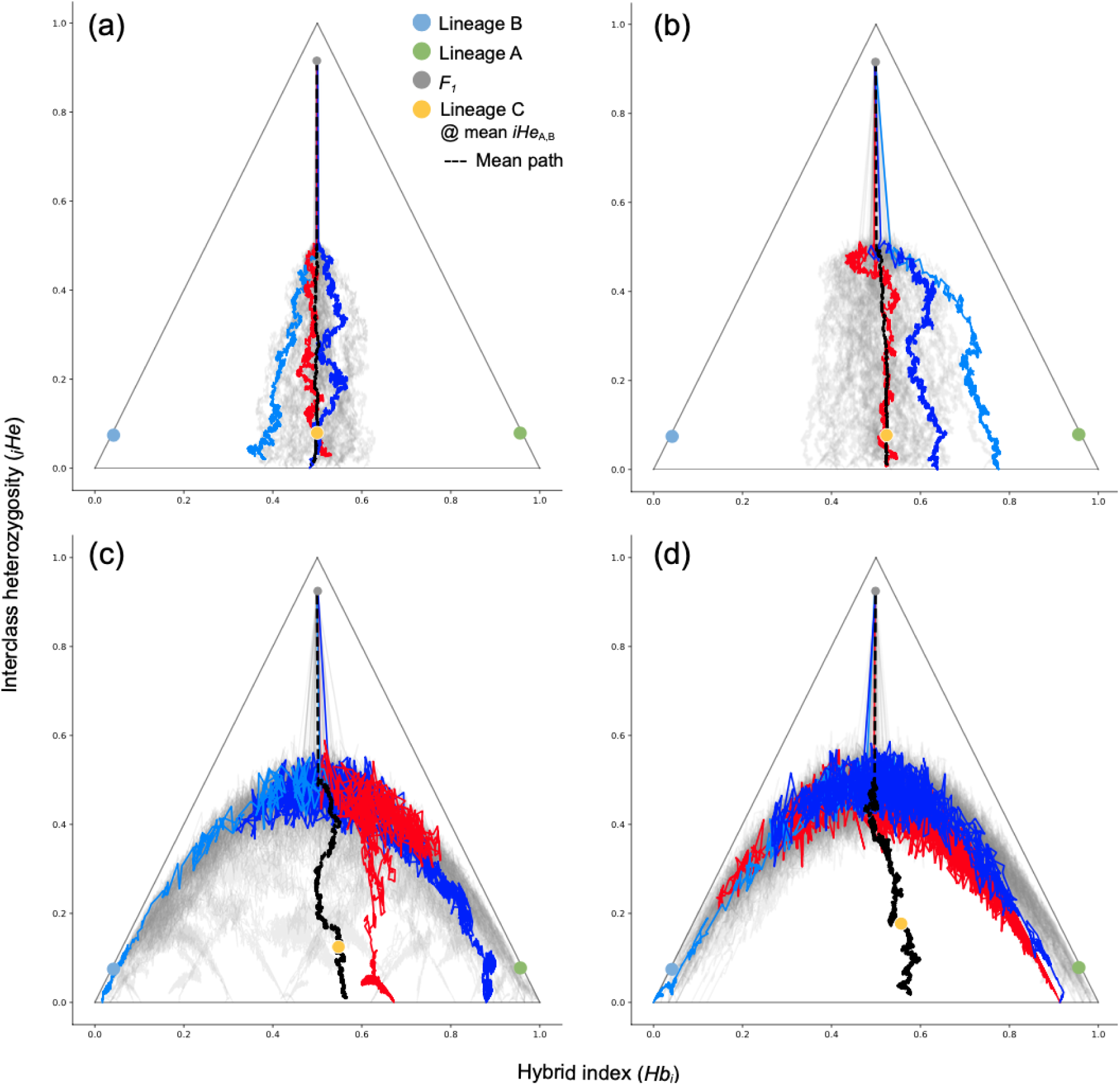
Triangle plots showing the joint distribution of ancestry (*Hb*_i_) and interclass heterozygosity (_i_*He*) for 50 replicate hybrid populations (lineage C) over 15 N_e_ generations under varying linkage scenarios: (a) free recombination, (b) single chromosome, (c) tight linkage (0.5 cM), and (d) complete linkage (no recombination). Grey lines show population trajectorie from the F₁ generation; mean lineage A (green circle), lineage B (blue circle), F₁ (grey circle), and mean of lineage C at the time of reaching mean parental _i_*He* (mean _i_*He*_AB_) (yellow circle). Highlighted trajectories (dark blue, light blue, red) represent a random path, an outlier, and the trajectory closest to the mean. The black dashed line shows the mean trajectory. Axes show mean hybrid index (x-axis) and mean interclass heterozygosity (y-axis).

## References

Barrera-Guzmán AO, Aleixo A, Shawkey MD, Weir JT. 2018. Hybrid speciation leads to novel male secondary sexual ornamentation of an Amazonian bird. Proceedings of the National Academy of Sciences of the United States of America 115: E218–E225.

Bateson W. 1909. Heredity and variation in modern lights. In: Seward AC (Ed.) Darwin and modern science. Cambridge University Press.

Blanckaert A, Bank C. 2018. In search of the Goldilocks zone for hybrid speciation. Plos Genetics 14: 23.

Blanckaert A, Sriram V, Bank C. 2023. In search of the Goldilocks zone for hybrid speciation II: Hard times for hybrid speciation? Evolution 77: 2162–2172.

Boca SM, Huang L, Rosenberg NA. 2020. On the heterozygosity of an admixed population. Journal of Mathematical Biology 81: 1217–1250.

Bryant D, Bouckaert R, Felsenstein J, Rosenberg NA, RoyChoudhury A. 2012. Inferring species trees directly from biallelic genetic markers: Bypassing gene trees in a full coalescent analysis. Molecular Biology and Evolution 29: 1917–1932.

Corneault AA, Matute DR. 2018. Genetic divergence and the number of hybridizing species affect the path to homoploid hybrid speciation. Proceedings of the National Academy of Sciences of the United States of America 115: 9761–9766.

Coyne JA, Orr HA. 2004. Speciation. Sinauer, Sunderland.

Dittrich-Reed DR, Fitzpatrick BM. 2013. Transgressive hybrids as hopeful monsters. Evolutionary Biology 40: 310–315.

Dobzhansky T. 1936. Studies on hybrid sterility. II. Localization of sterility factors in Drosophila pseudoobscura hybrids. Genetics 21: 113–135.

Durand EY, Patterson N, Reich D, Slatkin M. 2011. Testing for ancient admixture between closely related populations. Molecular Biology and Evolution 28: 2239–2252.

Earl DA, vonHoldt BM. 2012. STRUCTURE HARVESTER: A website and program for visualizing STRUCTURE output and implementing the Evanno method. Conservation Genetics Resources 4: 359–361.

Eaton DAR, Overcast I. 2020. IPYRAD: Interactive assembly and analysis of RADseq datasets. Bioinformatics 36: 2592–2594.

Elgvin TO, Trier CN, Torresen OK, Hagen IJ, Lien S, Nederbragt AJ, Ravinet M, Jensen H, Sætre GP. 2017. The genomic mosaicism of hybrid speciation. Science Advances 3: 15.

Emerson BC, Campedel A, Noguerales V. 2025. Island colonization and extinction of an insect species revealed by mitochondrial and genome-wide nuclear variation. Evolutionary Journal of the Linnean Society 4: kzaf012.

Eroukhmanoff F, Bailey RI, Sætre GP. 2013. Hybridization and genome evolution I: The role of contingency during hybrid speciation. Current Zoology 59: 667–674.

Evanno G, Regnaut S, Goudet J. 2005. Detecting the number of clusters of individuals using the software STRUCTURE: a simulation study. Molecular Ecology 14: 2611–2620.

Excoffier L, Marchi N, Marques DA, Matthey-Doret R, Gouy A, Sousa VC. 2021. FASTSIMCOAL2: Demographic inference under complex evolutionary scenarios. Bioinformatics 37: 4882–4885.

Excoffier L, Dupanloup I, Huerta-Sánchez E, Sousa VC, Foll M. 2013. Robust demographic inference from genomic and SNP data. Plos Genetics 9: e1003905.

Excoffier L, Foll M, Petit RJ. 2009. Genetic consequences of range expansions. Annual Review of Ecology Evolution and Systematics 40: 481–501.

Fitzpatrick BM. 2012. Estimating ancestry and heterozygosity of hybrids using molecular markers. BMC Evolutionary Biology 12:14.

García-Olivares V, López H, Patiño J, Alvarez N, Machado A, Carracedo JC, Soler V, Emerson BC. 2017. Evidence for mega-landslides as drivers of island colonization. Journal of Biogeography 44: 1053–1064.

Gompert Z, Fordyce JA, Forister ML, Shapiro AM, Nice CC. 2006. Homoploid hybrid speciation in an extreme habitat. Science 314: 1923–1925.

Grant V. 1971. Plant speciation. Columbia University Press, New York.

Gross BL, Rieseberg LH. 2005. The ecological genetics of homoploid hybrid speciation. Journal of Heredity 96: 241–252.

Hermansen JS, Sæther SA, Elgvin TO, Borge T, Hjelle E, Sætre GP. 2011. Hybrid speciation in sparrows I: Phenotypic intermediacy, genetic admixture and barriers to gene flow. Molecular Ecology 20: 3812–3822.

Hewitt GM. 2004. Genetic consequences of climatic oscillations in the Quaternary. Philosophical Transactions of the Royal Society of London Series B-Biological Sciences 359: 183–195.

Janes JK, Miller JM, Dupuis JR, Malenfant RM, Gorrell JC, Cullingham CI, Andrew RL. 2017. The *K*=2 conundrum. Molecular Ecology 26: 3594–3602.

Kadereit JW. 2015. The geography of hybrid speciation in plants. Taxon 64: 673–687.

Kagawa K, Takimoto G, Seehausen O. 2023. Transgressive segregation in mating traits drives hybrid speciation. Evolution 77: 1622–1633.

Kalyaanamoorthy S, Minh BQ, Wong TKF, von Haeseler A, Jermiin LS. 2017. MODELFINDER: Fast model selection for accurate phylogenetic estimates. Nature Methods 14: 587–589.

Katoh K, Standley DM. 2013. MAFFT - Multiple sequence alignment software version 7: improvements in performance and usability. Molecular Biology and Evolution 30: 772–780.

Lamichhaney S, Han F, Webster MT, Andersson L, Grant BR, Grant PR. 2018. Rapid hybrid speciation in Darwin’s finches. Science 359: 224–227.

Lewis PO. 2001. A likelihood approach to estimating phylogeny from discrete morphological character data. Systematic Biology 50: 913–925.

Linder CR, Rieseberg LH. 2004. Reconstructing patterns of reticulate evolution in plants. American Journal of Botany 91: 1700–1708.

Long ZQ, Rieseberg LH. 2024. Documenting homoploid hybrid speciation. Molecular Ecology 9: e17412.

Lopes F, Oliveira LR, Beux Y, Kessler A, Cárdenas-Alayza S, Majluf P, Páez-Rosas D, Chaves J, Crespo E, Brownell RL, Jr., et al. 2023. Genomic evidence for homoploid hybrid speciation in a marine mammal apex predator. Science Advances 9: 14.

Mavárez J, Salazar CA, Bermingham E, Salcedo C, Jiggins CD, Linares M. 2006. Speciation by hybridization in Heliconius butterflies. Nature 441: 868–871.

Melo MC, Salazar C, Jiggins CD, Linares M. 2009. Assortative mating preferences among hybrids offers a route to hybrid speciation. Evolution 63: 1660–1665.

Meyer A, Salzburger W, Schartl M. 2006. Hybrid origin of a swordtail species (Teleostei: Xiphophorus clemenciae) driven by sexual selection. Molecular Ecology 15: 721–730.

Mijangos JL, Gruber B, Berry O, Pacioni C, Georges A. 2022. dartR v2: An accessible genetic analysis platform for conservation, ecology and agriculture. Methods in Ecology and Evolution 13: 2150–2158.

Minh BQ, Nguyen MAT, von Haeseler A. 2013. Ultrafast approximation for phylogenetic bootstrap. Molecular Biology and Evolution 30: 1188–1195.

Muller HJ. 1942. Isolating mechanisms, evolution, and temperature. Biology Symposium 6: 71–125.

Muto N, Su Y-C, Hata H, Quan VN, Vilasri V, Ghaffar MA, Babaran RP. 2025. Homoploid hybrid speciation in a marine pelagic fish. Molecular Ecology, in press: e70112.

Nice CC, Gompert Z, Fordyce JA, Forister ML, Lucas LK, Buerkle CA. 2013. Hybrid speciation and independent evolution in lineages of alpine butterflies. Evolution 67: 1055–1068.

Nieto Feliner G, Álvarez I, Fuertes-Aguilar J, Heuertz M, Marques I, Moharrek F, Piñeiro R, Riina R, Rossello JA, Soltis PS, et al. 2017. Is homoploid hybrid speciation that rare? An empiricist’s view. Heredity 118: 513–516.

Noguerales V, Arjona Y, García-Olivares V, Machado A, López H, Patiño J, Emerson BC. 2024. Genetic legacies of mega-landslides: Cycles of isolation and contact across flank collapses in an oceanic island. Molecular Ecology 33: 13.

Noguerales V, Cordero PJ, Ortego J. 2016. Hierarchical genetic structure shaped by topography in a narrow-endemic montane grasshopper. BMC Evolutionary Biology 16: 96.

Noguerales V, Emerson BC. 2025. Arthropod mtDNA paraphyly: A case study of introgressive origin. Journal of Evolutionary Biology 38: 272–283.

Noguerales V, Ortego J. 2022. Genomic evidence of speciation by fusion in a recent radiation of grasshoppers. Evolution 76: 2618–2633.

Nolte AW, Freyhof J, Stemshorn KC, Tautz D. 2005. An invasive lineage of sculpins, Cottus sp. (Pisces, Teleostei) in the Rhine with new habitat adaptations has originated from hybridization between old phylogeographic groups. Proceedings of the Royal Society B-Biological Sciences 272: 2379–2387.

Olave M, Nater A, Kautt AF, Meyer A. 2022. Early stages of sympatric homoploid hybrid speciation in crater lake cichlid fishes. Nature Communications 13: 9.

Ortego J, Gutierrez-Rodríguez J, Noguerales V. 2021. Demographic consequences of dispersal-related trait shift in two recently diverged taxa of montane grasshoppers. Evolution 75: 1998–2013.

Ortego J, Noguerales V. 2025. Mountain speciation driven by high rates of lineage formation and rapid evolution of partial reproductive isolation: insights from a recent radiation of grasshoppers (Orthoptera: Gomphocerinae). Zoological Journal of the Linnean Society 205: zlaf141.

Pal A, Shipilina D, Le Moan A, McNairn AJ, Grenier JK, Kucka M, M Coop G, Chan YF, Barton NH, Field DL, Stankowski S. 2025. Genealogical analysis of replicate flower colour hybrid zones in Antirrhinum. Molecular Ecology, in press: e70067.

Pérez-Delgado AJ, Arribas P, Hernando C, López H, Arjona Y, Suárez-Ramos D, Emerson BC, Andújar C. 2022. Hidden island endemic species and their implications for cryptic speciation within soil arthropods. Journal of Biogeography 49: 1367–1380.

Peterson BK, Weber JN, Kay EH, Fisher HS, Hoekstra HE. 2012. Double digest RADseq: An inexpensive method for de novo SNP discovery and genotyping in model and non-model species. Plos One 7: e37135.

Pritchard JK, Stephens M, Donnelly P. 2000. Inference of population structure using multilocus genotype data. Genetics 155: 945–959.

R Core Team. 2022. R: A language and environment for statistical computing. R Foundation for Statistical Computing, Vienna. Available at https://www.r-project.org/.

Rohwer S, Martin PR. 2007. Time since contact and gene flow may explain variation in hybrid frequencies among three Dendroica townsendi x D. occidentalis (Parulidae) hybrid zones. Auk 124: 1347–1358.

Roux C, Fraïsse C, Romiguier J, Anciaux Y, Galtier N, Bierne N. 2016. Shedding light on the grey zone of speciation along a continuum of genomic divergence. Plos Biology 14: 22.

Runemark A, Trier CN, Eroukhmanoff F, Hermansen JS, Matschiner M, Ravinet M, Elgvin TO, Sætre GP. 2018. Variation and constraints in hybrid genome formation. Nature Ecology and Evolution 2: 549–556.

Runemark A, Vallejo-Marin M, Meier JI. 2019. Eukaryote hybrid genomes. Plos Genetics 15: 22.

Salces-Castellano A, Patiño J, Alvarez N, Andújar C, Arribas P, Braojos-Ruiz JJ, Del Arco-Aguilar M, Garcia-Olivares V, Karger DN, López H, et al. 2020. Climate drives community-wide divergence within species over a limited spatial scale: Evidence from an oceanic island. Ecology Letters 23: 305–315.

Salces-Castellano A, Stankowski S, Arribas P, Patiño J, Karger DN, Butlin R, Emerson BC. 2021. Long-term cloud forest response to climate warming revealed by insect speciation history. Evolution 75: 231–244.

Schumer M, Cui RF, Rosenthal GG, Andolfatto P. 2015. Reproductive isolation of hybrid populations driven by genetic incompatibilities. Plos Genetics 11: 21.

Schumer M, Rosenthal GG, Andolfatto P. 2014. How common is homoploid hybrid speciation? Evolution 68: 1553–1560.

Schumer M, Rosenthal GG, Andolfatto P. 2018. What do we mean when we talk about hybrid speciation? Heredity 120: 379–382.

Schwarz D, Matta BM, Shakir-Botteri NL, McPheron BA. 2005. Host shift to an invasive plant triggers rapid animal hybrid speciation. Nature 436: 546–549.

Schwarzbach AE, Donovan LA, Rieseberg LH. 2001. Transgressive character expression in a hybrid sunflower species. American Journal of Botany 88: 270–277.

Sobel JM, Chen GF. 2014. Unification of methods for estimating the strength of reproductive isolation. Evolution 68: 1511–1522.

Stamatakis A. 2014. RAXML version 8: A tool for phylogenetic analysis and post-analysis of large phylogenies. Bioinformatics 30: 1312–1313.

Stüben PE, Behne L, Floren A, Günther H, Klopfstein S, López H, Machado A, Schwarz M, Wägele JW, Wunderlich J, Astrin JJ. 2010. Canopy fogging in the Canarian laurel forest of Tenerife and La Gomera. Weevil News 51: 21.

Stüben PE. 2022. Weevils of Macaronesia: Canary Islands, Madeira, Azores (Coleoptera: Curculionoidea). Curculio Institute, Mönchengladbach, Germany.

Trifinopoulos J, Nguyen LT, von Haeseler A, Minh BQ. 2016. W-IQ-TREE: A fast online phylogenetic tool for maximum likelihood analysis. Nucleic Acids Research 44: W232–W235.

Wang D, Xu X, Zhang H, Xi Z, Abbott RJ, Fu J, Liu J. 2022. Abiotic niche divergence of hybrid species from their progenitors. The American Naturalist 200: 634–645.

Wang ZF, Kang MH, Li JL, Zhang ZY, Wang YF, Chen CL, Yang YZ, Liu JQ. 2022. Genomic evidence for homoploid hybrid speciation between ancestors of two different genera. Nature Communications 13: 9.

Wiens B, Decicco LH, Colella JP. 2025. triangulaR: An R package for identifying AIMs and building triangle plots using SNP data from hybrid zones. Heredity 134: 251–262.

Yu DW, Ji Y, Emerson BC, Wang X, Ye C, Yang C, Ding Z. 2012. Biodiversity soup: metabarcoding of arthropods for rapid biodiversity assessment and biomonitoring. Methods in Ecology and Evolution 3: 613–623.

Zhao DD, Zhang JG, Hui N, Wang L, Tian Y, Ni WN, Long JH, Jiang L, Li Y, Diao SF, et al. 2023. A genomic quantitative study on the contribution of the ancestral-state bases relative to derived bases in the divergence and local adaptation of Populus davidiana. Genes 14: 22.

Zou T, Kuang W, Yin T, Frantz L, Zhang C, Liu J, Wu H, Yu L. 2022. Uncovering the enigmatic evolution of bears in greater depth: The hybrid origin of the Asiatic black bear. Proceedings of the National Academy of Sciences of the United States of America. 119: e2120307119.

## References

Andrews S. 2010. FASTQC: A quality control tool for high throughput sequence data. http://www.bioinformatics.babraham.ac.uk/projects/fastqc/

Bouckaert R, Heled J, Kuehnert D, Vaughan T, Wu C-H, Xie D, Suchard MA, Rambaut A, Drummond AJ. 2014. BEAST2: A software platform for Bayesian evolutionary analysis. PLOS Computational Biology 10: e1003537.

Bouckaert RR. 2010. DENSITREE: Making sense of sets of phylogenetic trees. Bioinformatics 26: 1372–1373.

Burnham KP, Anderson DR. 2002. Model selection and multimodel inference. A practical information-theoretic approach. 2^nd^ ed. Springer-Verlag, New York.

de Manuel M, Kuhlwilm M, Frandsen P, Sousa VC, Desai T, Prado-Martinez J, Hernandez-Rodriguez J, Dupanloup I, Lao O, Hallast P, Schmidt JM, Heredia-Genestar JM, Benazzo A, Barbujani G, Peter BM, Kuderna LFK, Casals F, Angedakin S, Arandjelovic M, Boesch C, Kuehl H, Vigilant L, Langergraber K, Novembre J, Gut M, Gut I, Navarro A, Carlsen F, Andres AM., Siegismund HR, Scally A, Excoffier L, Tyler-Smith C, Castellano S, Xue Y, Hvilsom C, Marques-Bonet T. 2016. Chimpanzee genomic diversity reveals ancient admixture with bonobos. Science 354: 477–481.

Durand EY, Patterson N, Reich D, Slatkin M. 2011. Testing for ancient admixture between closely related populations. Molecular Biology and Evolution 28: 2239–52.

Drummond AJ, Rambaut A. 2007. BEAST: Bayesian evolutionary analysis by sampling trees. BMC Evolutionary Biology 7: 214.

Emerson BC, Casquet J, Lopez H, Cardoso P, Borges PAV, Mollaret N, Oromi P, Strasberg D, Thebaud C. 2017. A combined field survey and molecular identification protocol for comparing forest arthropod biodiversity across spatial scales. Molecular Ecology Resources 17: 694–707.

Evanno G, Regnaut S, Goudet J. 2005. Detecting the number of clusters of individuals using the software STRUCTURE: A simulation study. Molecular Ecology 14: 2611–2620.

Excoffier L, Marchi N, Marques DA, Matthey-Doret R, Gouy A, Sousa VC, Schwartz R. 2021. FASTSIMCOAL2: Demographic inference under complex evolutionary scenarios. Bioinformatics 37: 4882–4885.

Gilbert KJ, Andrew RL, Bock DG, Franklin MT, Kane NC, Moore J-S, Moyers BT, Renaut S, Rennison DJ, Veen T, Vines TH. 2012. Recommendations for utilizing and reporting population genetic analyses: The reproducibility of genetic clustering using the program STRUCTURE. Molecular Ecology 21: 4925–4930.

Jakobsson M, Rosenberg NA. 2007. CLUMPP: A cluster matching and permutation program for dealing with label switching and multimodality in analysis of population structure. Bioinformatics 23: 1801–1806.

Keightley PD, Ness RW, Halligan DL, Haddrill PR. 2014. Estimation of the spontaneous mutation rate per nucleotide site in a Drosophila melanogaster full-sib family. Genetics 196: 313–320.

Papadopoulou A, Knowles LL. 2015. Genomic tests of the species-pump hypothesis: Recent island connectivity cycles drive population divergence but not speciation in Caribbean crickets across the Virgin Islands. Evolution 69: 1501–1517.

Rambaut A, Drummond AJ, Xie D, Baele G, Suchard MA. 2018. Posterior summarization in Bayesian phylogenetics using TRACER 1.7. Systematic Biology 67: 901–904.

Rozas J, Ferrer-Mata A, Sánchez-del Barrio JC, Guirao-Rico S, Librado P, Ramos-Onsins SE, Sánchez-Gracia A. 2017. DNASP 6: DNA sequence polymorphism analysis of large data sets. Molecular Biology and Evolution 34: 3299–3302.

Salces-Castellano A, Patiño J, Alvarez N, Andújar C, Arribas P, Braojos-Ruiz JJ, del Arco-Aguilar M, García-Olivares V, Karger DN, López H, Manolopoulou I, Oromí P, Pérez-Delgado AJ, Peterman WE, Rijsdijk KF, Emerson BC. 2020. Climate drives community-wide divergence within species over a limited spatial scale: evidence from an oceanic island. Ecology Letters 23: 305–315.

Stamatakis A. 2014. RAXML version 8: A tool for phylogenetic analysis and post-analysis of large phylogenies. Bioinformatics, 30: 1312–1313.

Thomé MTC, Carstens BC. 2016. Phylogeographic model selection leads to insight into the evolutionary history of four-eyed frogs. Proceedings of the National Academy of Sciences of the United States of America 113: 8010–8017.

